# Angiopoietin 1 and integrin beta 1b are vital for zebrafish brain development

**DOI:** 10.1101/2021.04.14.439926

**Authors:** Yu-Chia Chen, Tomás A. Martins, Valentina Marchica, Pertti Panula

## Abstract

This study aimed at identifying the role of angiopoietin 1 (angpt1) in brain development, the mode of action of angpt1, and the main targets in the zebrafish brain. We investigated embryonic brain angiogenesis and neural development in the *angpt1*^*sa14264*^, *itgb1b*^*mi371*^, *tek*^*hu1667*^ mutant fish, and the effects of transgenic overexpression of *angpt1* in the larval brain. Lack of *angpt1* was associated with downregulation of *tek* and upregulation of *itgb1b*. We found deficiencies in the patterning of proliferation, the vascular network and reticulospinal neurons in the hindbrain, and selective deficiencies in specific neurotransmitter systems. In the *angpt1*^*sa14264*^ and *itgb1b*^*mi371*^ larval brains, using microangiography, retrograde labeling, and immunostaining, we demonstrated that the targeted destruction of *angpt1*^*sa14264*^ and *itgb1b*^*mi371*^ mutant fish caused severe irregular cerebrovascular development, aberrant hindbrain patterning, downregulation of neural proliferation, expansion of the radial glial progenitors, deficiencies of dopaminergic, histaminergic, and GABAergic populations in the larval brain. In contrast, the *tek*^*hu1667*^ mutants regularly grew with no such apparent phenotypes. Neurally overexpressed *angpt1* promoted opposite effects by increasing the vascular branching, increasing cell proliferation, and neuronal progenitors. Notably, zebrafish *angpt1* showed neurogenic activity independent of its typical receptor *tek*, indicating the novel role of a dual regulation by *angpt1* in embryonic neurogenesis and angiogenesis in zebrafish. The results show that angpt1 and its interaction with itgb1b are crucial in zebrafish brain neuronal and vascular development and suggest that angpt1 through itgb1b can act as a neurogenic factor in the neural proliferation fate in the developing brain.

## Introduction

A functional neural orchestra requires reciprocal molecular communications between neurogenesis and angiogenesis in the nervous system (Walchli et al., 2015). The nervous and vascular networks develop in a similar stereotypic pattern and navigate in close proximity toward their targets as needed (Carmeliet and Tessier-Lavigne, 2005). Besides anatomical similarities, during embryonic development, these systems share common guiding cues and signaling pathways to establish the accurate nerve-vessel coordination such as slits and their receptor roundabouts, netrins and UNC5B, semaphorins and neuropilins, ephrins and ephr receptors, Wnts and Notch signaling pathways (Adams and Eichmann, 2010; Thomas et al., 2013). This mutual interdependency is remarkably conserved among vertebrates, including zebrafish (Melani and Weinstein, 2010). During embryonic neurogenesis and post-injury restoration, angiogenic factors can also act as neurotrophic factors to regulate nervous proliferation, differentiation, and neuronal remodeling in stem cell niches (Hatakeyama et al., 2020). For instance, vascular endothelial growth factors (VEGFs) via Notch signaling pathways play essential roles in promoting angiogenesis and vascular formation as well as proliferation and neurogenesis in adult stem cells (Lin et al., 2019). Likewise, angiopoietins including Angpt1, Angpt2, and Angpt3 are vital factors for cardiovascular development and adult vessel regeneration. Angiopoietin 1 (Angpt1) primarily binds to its endothelial receptor tyrosine kinase (Tek/ Tie-2) and activates the downstream PI3K/AKT and Rho family GTPase to support vessel development and stabilize the vasculature (Gale and Yancopoulos, 1999; Koh, 2013). Besides its crucial role in angiogenesis, growing evidence indicates that Angpt1 can induce neuronal differentiation and neurite outgrowth in ex vivo systems (Bai et al., 2009; Chen et al., 2009a; Rosa et al., 2010). Angpt1 acting as a neurotrophic factor, prevents blood-brain barrier (BBB) leakage in the ischemic rats through the increase of vascular density and neuronal differentiation (Meng et al., 2014). Previous studies have suggested that the Angpt1/tek pathway is not important in zebrafish in contrast to its crucial role in mammals (Gjini et al., 2011). Nevertheless, angpt1 appears to be an important regulator of brain size in guppies and zebrafish (Chen et al., 2015), and notch1 is crucial for this regulation. In this study, we explored the possibility that angpt1 is important in brain development, and instead of *tek*, *angpt1* could use another signaling system to regulate the development of both vasculature and neuronal development in the zebrafish brain. Previous studies have shown that fiber outgrowth induced by angpt1 in neuronal PC12 cells is independent of tek and dependent on beta 1 integrin (Chen et al., 2009a), and cell adhesion, not limited to endothelial cells, *angpt1* is not reliant on tek, but requires integrins (Carlson et al., 2001). To investigate the function of *angpt1* and its interaction with *tek* and *β1-integrin* (*itgb1b*) in neurogenesis, we used loss-of - function and gain-of-function approaches. We first examined the overt vasculature and neural phenotype of the *angpt1*^*sa14264*^ mutant zebrafish. Besides the *angpt1*^*sa14264*^ mutant, we further performed a comprehensive analysis of the neural development in the *itgb1b*^*mi371*^, *tek*^*hu1667*^ mutant lines, and transgenic overexpression of zebrafish *angpt1* larvae driven by ubiquitous and neural promoters to reveal the neurogenic effects in the zebrafish larval brain. Altogether, our findings suggest that angpt1 and itgb1b have vital dual functions in embryonic neurogenesis, notably, in a tek-independent manner in zebrafish.

## Materials and methods

### Zebrafish strain and maintenance

The zebrafish Turku wild-type (WT) strain was obtained from our breeding line maintained in the laboratory for more than two decades (Chen et al., 2020; Kaslin and Panula, 2001). The fish were raised at 28°C and staged in hours post-fertilization (hpf) or days post-fertilization (dpf) as described previously (Kimmel 1995). The *angpt1* mutant allele was generated by the Sanger Centre Zebrafish Project (The Wellcome Trust Sanger Institute, Hinxton, Cambridge, UK). The F3 heterozygous *angpt1*^*sa14264/+(TL)*^ strain was obtained from the European Zebrafish Resource Center and outcrossed with the Turku WT strain at least twice in our laboratory. All experiments were done using F6 or F7 progeny from in-crossed F5 or F6 *angpt1*^*sa14264/+*^ parents. The *itgb1b*^*mi371*^ mutant line (Iida et al., 2018) done by an N-ethyl-N-nitrosourea mutagenesis screen was provided by the National BioResource Project Zebrafish (NBRP; https://shigen.nig.ac.jp/zebra/index_en.html). The *tek*^*hu1667*^ mutant line was kindly provided by Dr. Stefan Schulte-Merker (Gjini et al., 2011). The experimental procedures in this study were conducted under the Office of the Regional Government of Southern Finland’s permits, in agreement with the European Convention’s ethical guidelines.

### Fin clipping and genomic DNA extraction of larval and adult zebrafish

To prepare genomic DNA for lysis, individual tail clippings were incubated in 50 μl of lysis buffer (10mM Tris-HCl pH8.3, 50mM KCl, 0.3%Tween 20 and 0.3%NP40) at 98°C for 10mim and then transferred on ice for 2 min followed by 1μl of Proteinase K (20mg/ml) digestion to remove proteins and then incubated at 55°C for at least 4 hours. To inactivate Proteinase K, the samples were set at 98°C for 10min and quenched on ice. The high-resolution melting (HRM) curve acquisition and analysis were used to detect the point mutation. Primers flanking the mutation site were designed using Primer-BLAST (https://www.ncbi.nlm.nih.gov/tools/primer-blast): forward_5’ AGAGCTACCGGAAACAGCAC and reverse_5’ GCGTCTTTAGCACAGAGGCT. The HRM analysis was done on the LightCycler 480 instrument (Roche), and reaction mixtures were: 1X LightCycler 480 HRM master mix (Roche), 2mM MgCl2, and 0,15μM primer mixtures. The PCR cycling protocol was: one cycle of 95°C for 10min; 45 cycles of 95°C for 10s, 60°C for 15s and 72°C for 20s followed by Melting curve acquisition: one cycle of 95°C for 60s and 40°C for 60s. PCR products were denatured at 95°C for 60s, renatured at 40°C for 60s, and melted at 60°c to 95°C with 25 signal acquisitions per degree. Melting curves were generated over a 65-95°C range. Curves were analyzed using the LightCycler 480 gene-scanning software (Roche) (Chen et al., 2020). To identify deviations of the curves indicating sequence polymorphism, a three-step analysis by the Gene Scanning program (Roche) was as followed (1) the first step was to normalize the raw melting-curve data by setting the initial fluorescence uniformly to a relative value of 100% while the final fluorescence to a relative value of 0%. (2) the second step was to determine the temperature threshold where the entire double-stranded DNA was completed denatured. (3) the final step was to further analyze the differences in melting-curve shapes to cluster samples into groups. Those showing similar melting curves were characterized as the same genotype. In this study, all embryos older than 3 dpf of *angpt1*^*sa14264*^, *itgb1b*^*mi371*^, *tek*^*hu1667*^ mutant lines were genotyped. 36-hpf and 48-hpf *angpt1*^*−/−*^ larvae (*angpt1 knockout (KO*)) were identified primarily by distinctly deficient cardiovascular circulation and then confirmed using genotyping. Because the phenotypic trait faithfully correlated with the genotyping outcome, the rest of the siblings with the regular cardiovascular system were marked as the *angpt1* WT without genotyping. Correctness of this procedure was controlled by genotyping samples of embryos.

### RNA isolation and cDNA synthesis

For quantitative real-time PCR analysis, total RNA was extracted from 30 pooled embryos or ten pooled genotyped larvae for each sample using the RNeasy mini Kit (Qiagen, Valencia, CA, USA). According to instructions provided by the manufacturer, to synthesize cDNA, one or two μg of total RNA was reverse-transcribed using the SuperScriptTM III reverse transcriptase (Invitrogen, Carlsbad, CA, USA).

### Quantitative real-time PCR (qPCR)

qPCR was performed using the Lightcycler® 480 SYBR Green I Master (Roche) in the LightCycler 480 instrument (Roche, Mannheim, Germany). Primers (Table 1) for amplification were designed by Primer-BLAST (NCBI). For developmental qPCR, *β-actin* and *lsm12b* were used as reference controls; for the 3-dpf qPCR analysis, *β-actin* and *ribosomal protein L13a* (*rpl13a*) were used to standardize the results (Hu et al., 2016). All primer sets’ specificity was confirmed to amplify only a single product of the correct size by melting curve analysis, and no peaks appeared when RNA samples without reverse transcriptase was added. Cycling parameters were as follows: 95°C for 5 min and 45 cycles of the following, 95°C for 10s, 60°C for 15s and 72°C for 20s. Fluorescence changes were monitored after each cycle. Dissociation curve analysis was performed (0.1°C per increase from 60°C to 95°C with continuous fluorescence readings) at the end of the cycles to ensure that only a single amplicon was obtained. All reactions were performed in technical duplicates, and data are shown in biological triplicates. Results were analyzed with the LightCycler 480 software with the default settings. The comparative C_T_ method (2^−ΔCT^) calculated relative quantitative gene expressions by comparing C_T_ values of the gene of interest relative to internal control genes (Schmittgen and Livak, 2008). Since the gene expression changes showed similar trends when normalized to different housekeeping genes (data not shown), the results normalized to *β-actin* or *lsm12b* are shown in this study.

**Table 1.**
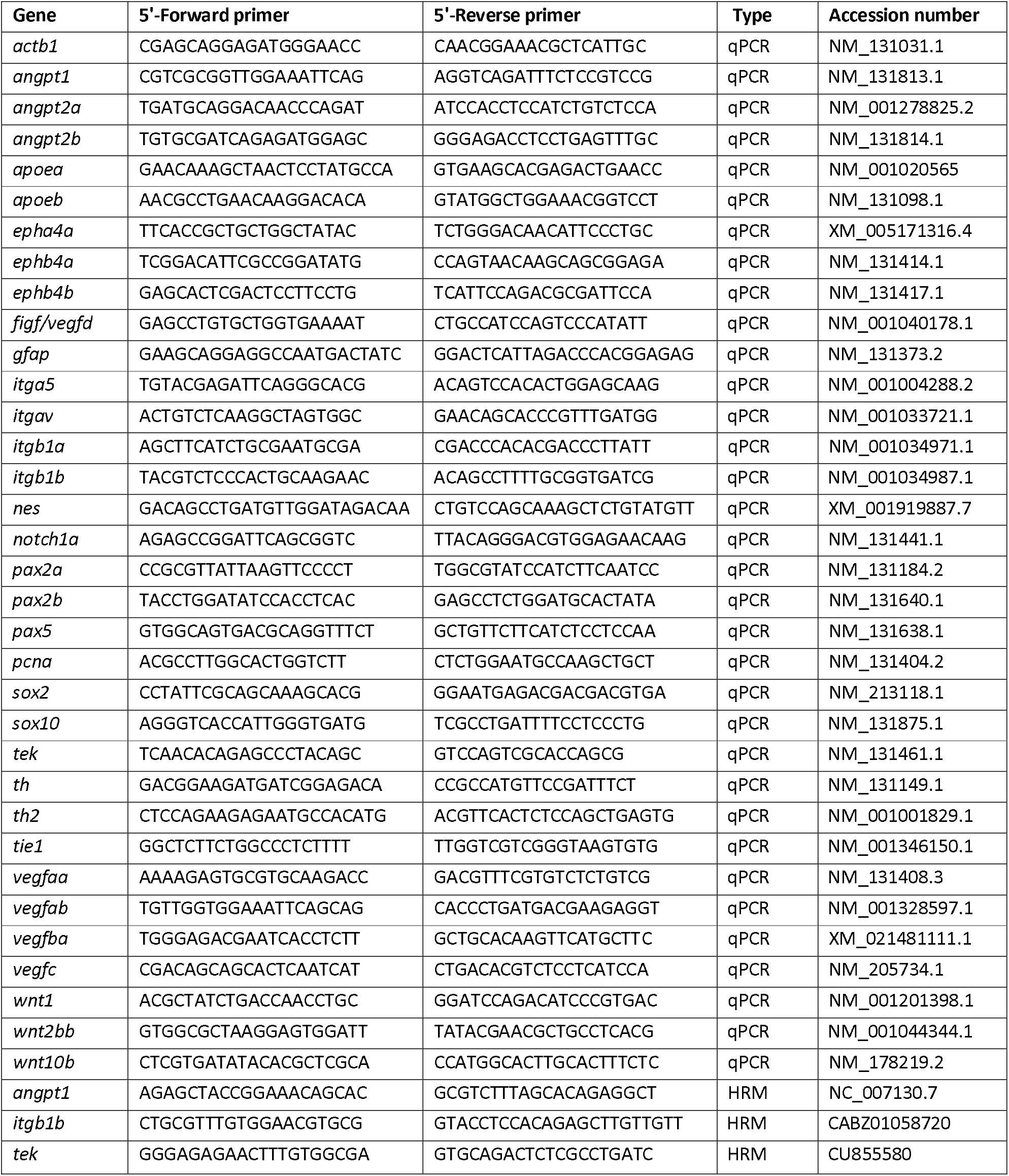
List of primers used in this study.

### Fixation of embryos

To generate transparent zebrafish for *in situ* hybridization, Turku WT embryos were treated with 0.03% 1-phenyl-2-thiourea (PTU) in E3 medium (5mM NaCl, 0.17mM KCl, 0,4 mM CaCl2, and 0.16mM MgSO4) from 13 hpf to 3 dpf to prevent pigmentation and the PTU-treated embryos were fixed in 4% paraformaldehyde (PFA) in PBS overnight at 4°C. The *angpt1* KO and sibling WT embryos were sensitive to PTU treatment, so the *angpt1* embryos were fixed in 4% PFA in PBS for three h and then treated with 3%H_2_O_2_/ 0.5 % KOH in PBS for 30 min at 25°C (Thisse and Thisse, 2008) and washed several times with PBS to remove pigmentations. The fixed embryos were dehydrated with a graded methanol series (25%, 50%, and 75% for 10min each) and stored at −20°C in 100% methanol. Samples for Immunostaining were fixed in 2% PFA or 4% 1-ethyl-3 (3-dimethylaminopropyl)-carbodiimide (EDAC, Carbosynth, Berkshire, UK). The fixed brains were dissected to enhance antigen presentation and to improve image quality.

### Whole-mount *in situ* hybridization (WISH)

Whole-mount *in situ* hybridization (WISH) was performed on 4% PFA fixed embryos as described previously (Chen et al., 2009b). Briefly, antisense and sense digoxogenin (DIG)-labeled RNA probes were generated using the DIG RNA labeling kit (Roche Diagnostics, Germany), following the manufacturer’s instructions. The WISH procedure followed Thisse’s protocol (Thisse and Thisse, 2008). The specificity of antisense riboprobes was determined by using sense probes showing faint or no staining signals. The pre-hybridization and hybridization were conducted at 65°C. *In situ* hybridization signals were detected with sheep anti-digoxigenin-AP Fab fragments (1:10000; Roche Diagnostics, Germany) conjugated with alkaline phosphatase. The colorimetric staining was carried out with chromogen substrates (nitro blue tetrazolium and 5-bromo-4-chloro-3-indolil-phosphate).

### Heart and blood circulation rate measurement

2-dpf *angpt1* KO and WT embryos were anesthetized with 0.0168% (w/v) tricaine (MS-222, Sigma) and mounted on 3% methylcellulose (Sigma, M0262) for imaging. Heart and blood circulation videos were recorded using an Olympus IX70 microscope (Olympus, Tokyo, Japan) and a Hamamatsu ORCA-Flash 4.0 CMOS camera with HCImage software (Hamamatsu Photonics, Shizuoka, Japan). The recording speed was 50 fps or 133 fps for heart beats and blood circulation, respectively. Videos were analyzed frame by frame with ImageJ. Quantification of the heart rate was done using manual counting by a blinded tester, and the blood circulation velocity was determined by tracking the flow of red blood cells based on Paavola et al. (Paavola et al., 2013).

### Microangiography

Microangiography was performed on 3-dpf *angpt1* and *itgb1b* mutant embryos. Anesthetized embryos were placed on a 1% agarose injection stage. Approximately 1.5 nl of solution containing 2mg/ml of fluorescein isothiocyanate-dextran 2000kDa (FITC-Dextran 2000, Sigma 52471) was injected into the sinus venosa of anesthetized embryos (Schmitt et al., 2012). Injected embryos were embedded in 2% low-melting agarose and live imaged by confocal microscopy within 20 min after injections.

### EdU labeling and double Immunocytochemistry

The Click-iT^™^EdU Alexa Fluor 488 imaging kit (Molecular Probes) was used according to the manufacturer’s instruction with minor modifications to detect S-phase proliferation of dividing cells. Briefly, 5-dpf larvae were incubated in 0.5mM EdU/ E3 buffer with 1%DMSO for 24h at 28°C. EdU-labeled samples were transferred back to E3 for 30 min and fixed in 2% PFA overnight at 4°C with agitations (Chen et al., 2020). The specimens were dissected to enhance sample penetration. Dissected brains were incubated with monoclonal mouse anti-HuC/D (1:500, Invitrogen, Cat.No: A21271), anti-tyrosine hydroxylase (TH1) monoclonal mouse antibody (1:1000; Product No 22941, Immunostar, Husdon, WI, USA), anti-GABA 1H (1:1000; (Karhunen et al., 1993; Kukko-Lukjanov and Panula, 2003)and mouse anti-zrf1 (Gfap; 1:1000, Zebrafish International Resource Center). The specificities of the anti-GABA and anti-histamine, commercial anti- mouse monoclonal TH, antibodies have been verified previously (Kaslin and Panula, 2001). The following secondary antibodies were applied: Alexa Fluor^®^ 488 and 647 anti-mouse or anti-rabbit IgG (1:1000; Invitrogen, Eugene, OR, USA). After immunostaining, labeled specimens were fixed in 4% PFA for 20 min at RT and then Incubated in a 1XClick-iT EdU cocktail containing Alexa488-azide for one hour in the dark at room temperature. After removal of the reaction cocktail and rinsing 3 times in 1XPBST for 10min, the samples were mounted in 80% glycerol/ PBS for confocal microscopy.

### Hemoglobin staining

Dechorionated embryos were incubated in o-dianisidine solution (0.6mg/ml of o-dianisidine; D1943, Sigma; 10 mM sodium acetate pH4.5; 0.65% H_2_O_2_; 40% ethanol) in the dark for 30 min at RT. The stained embryos were washed with ddH_2_O 3 times for 10 min and fixed in 4% PFA for two hours at RT. Pigments were removed by incubation in the bleaching solution containing 0.8% KOH, 0.9% H_2_O_2,_ and 0.1% Tween-20. The stained samples were washed with 1xPBST 4 times for 15 min and embedded in 50% glycerol/ PBS for microscopy (Huang et al., 2014).

### Retrograde labeling

Reticulospinal neurons were visualized by retrograde labeling with the fluorescent dye, Dextran Alexa Flour 488 10,000 KDa (Invitrogen D22910) applied to the spinal cord of 3-dpf larvae (Alexandre et al., 1996). Briefly, anesthetized larvae were placed on a 1% agarose template. The labeling dye was applied for 2 min on a spinal cord lesion site between somite 12 and 15 made with a 30G needle in the dark, and injected embryos were incubated in E3 medium for two hours. The labeled embryos were then killed on ice and fixed in 4% PFA. The fixed specimens were bleached (0.8%KOH, 0.9%H_2_O_2_, and 0.1%Tween-20) to avoid pigment disturbance while imaging and mounted in 80% glycerol for confocal microscopy.

### Imaging

Bright-field images were taken with a Leica DM IL inverted microscope. A Leica DM IRB inverted microscope with DFC 480 charge-coupled device camera was used to collect z-stacks of images which were processed with Leica Application Suite software. Immunofluorescence samples were examined using a Leica TCS SP2 AOBS confocal microscope. For excitation, an Argon laser (488 nm), green diode laser (561 nm) and red HeNe laser (633nm) were used. Emission was detected at 500-550 nm, 560-620 nm, and 630-680 nm, respectively. Crosstalk between the channels and background noise was eliminated with sequential scanning and frame averaging as described earlier (Sallinen et al., 2009). Stacks of images taken at 0.2 - 1.2 μm intervals were compiled, and the maximum intensity projection algorithm was used to produce final images with Leica Confocal software and Imaris imaging software version 6.0 (Bitplane AG, Zurich, Switzerland).

### Construction of ectopic angpt1 expression plasmids

*Angpt1* plasmids for ectopic expression were constructed by the Tol2kit based on Kwan et al. (Kwan 2007). 5’-entry vector pENTR5’-ubi (ubiquitin promoter) (Addgene plasmid #27230, p5E-h2afx from Tol2Kit, p5E-elavl3 (plasmid #72640), and p5E-gfap (plasmid #75024) were obtained from Addgene (Don et al., 2017; Mosimann et al., 2011). The intention was to investigate the role of ubiquitous, neuronal and non-neuronal expression of *angpt1* on neuronal markers and differentiation, respectively. The full-length coding sequence of zebrafish *angpt1* (NM_131813) was verified and reported in Chen et al. (Chen et al., 2015). The open reading frame *angpt1* was added to *HindIII* and *EcoRI* sites and ligated to the middle-entry pME-MCS vector. Gateway entry vectors (pENTR5’-promoter and p3E-EGFPpA) and destination vector pDestTol2CG2 were assembled using Gateway LR clonase II Mix (ThermoFisher, 11791-020), and Tol2 transposase sites flanked the entire cassette. The Tol2 pCS-TP plasmid kindly provided by Dr. Koichi Kawakami (Kawakami et al., 2004) was used to synthesize mRNA encoding Tol2 transposase using anSP6 mMESSAGE mMACHINE kit (ambion). Transgene expression in zebrafish was generated using Tol2-mediated transgenesis: the injection volume was 2 nl. 20ng/μl of ectopic *angpt1* expression plasmid combined with 50ng/μl of *Tol2* mRNA were injected into one-cell stage eggs. The control group was injected with 20ng/μl of the destination vector pDestTol2CG2 with 50ng/μl *Tol2* mRNA. Injected embryos with a reporter fluorescence marker in the heart were collected for further experiments.

### Statistical analysis

Data analysis was performed by GraphPad Prism v.7.0c software (San Diego, CA, USA). P-values were generated by one-way analysis of variance (ANOVA) for multiple comparisons using Dunnett’s test and Student’s t-test (unpaired test) to compare two groups. Data were presented as mean +/− SEM. P-value <0.05 was considered statistically significant.

## Results

### Spatiotemporal expression of angiogenic factors in developing embryos

Zebrafish neurulation starts at the 80% epiboly (end of gastrulation) stage (Lowery 2004). To gain insight into the roles of angiogenic factors in early embryonic neurogenesis, we first studied the spatiotemporal distribution of angiogenic factors during zebrafish embryogenesis. We conducted an extensive characterization of expression patterns of *angpt1, angpt2a, angpt2b, tie1*, and *tek* (*tie2*) from the zygotic period (0 hpf) to the hatching period (72 hpf). Among these angiogenic factors, by WISH, *angpt1* and its receptor *tek* were first detected in the yolk and the yolk syncytial layer at the one-cell stage (Fig. 1A). All angiopoietins and their receptors showed a ubiquitous distribution in the ectoderm and mesoderm from the blastula to the gastrula period (Fig. 1B, 1C), of which *angpt1* and *tek* displayed the most intense signals at early segmentation period (bud) (Fig. 1C). From the prim-5 stage, *angpt1* and *tek* were restrictedly present in the midbrain (Fig. 1D), eyes, and posterior cardinal channels (Fig.1E). The other factors were mainly found in the head (Fig. 1D). Throughout the long-pec period, all transcripts were primarily detected in the vasculature in the head including the otic placode, trabecula crani, and lower jaw and the trunk (Fig. 1F). A faint expression of *angpt1, tie1*, and *tek* was observed in the brain (Fig. 1F, 1G). To further distinguish each angiopoietin’s differential expression profiles, qPCR was performed on pooled wild-type whole larvae from 7 different developmental stages (Fig. 1H, 1I). The *angpt1* displayed the relatively highest maternal RNA expression compared with other angiogenic factors, and a maternal-to-zygotic transition change appeared at five hpf (Fig. 1I). Remarkably, the mRNA level of *angpt1, angpt2a, tie1*, and *tek* had reached a peak at the prim-5 stage and remained relatively stable over time afterwards; in contrast, the *angpt2a* mRNA persisted at a low-level expression during embryogenesis in agreement with the WISH results at all stages tested (Fig. 1H). By comparison, the *angpt1, tie1, and tek* revealed more intense expression during early development, suggesting the potential impact of *angpt1* and its associated receptors from the very early embryogenesis.

**Fig. 1.**
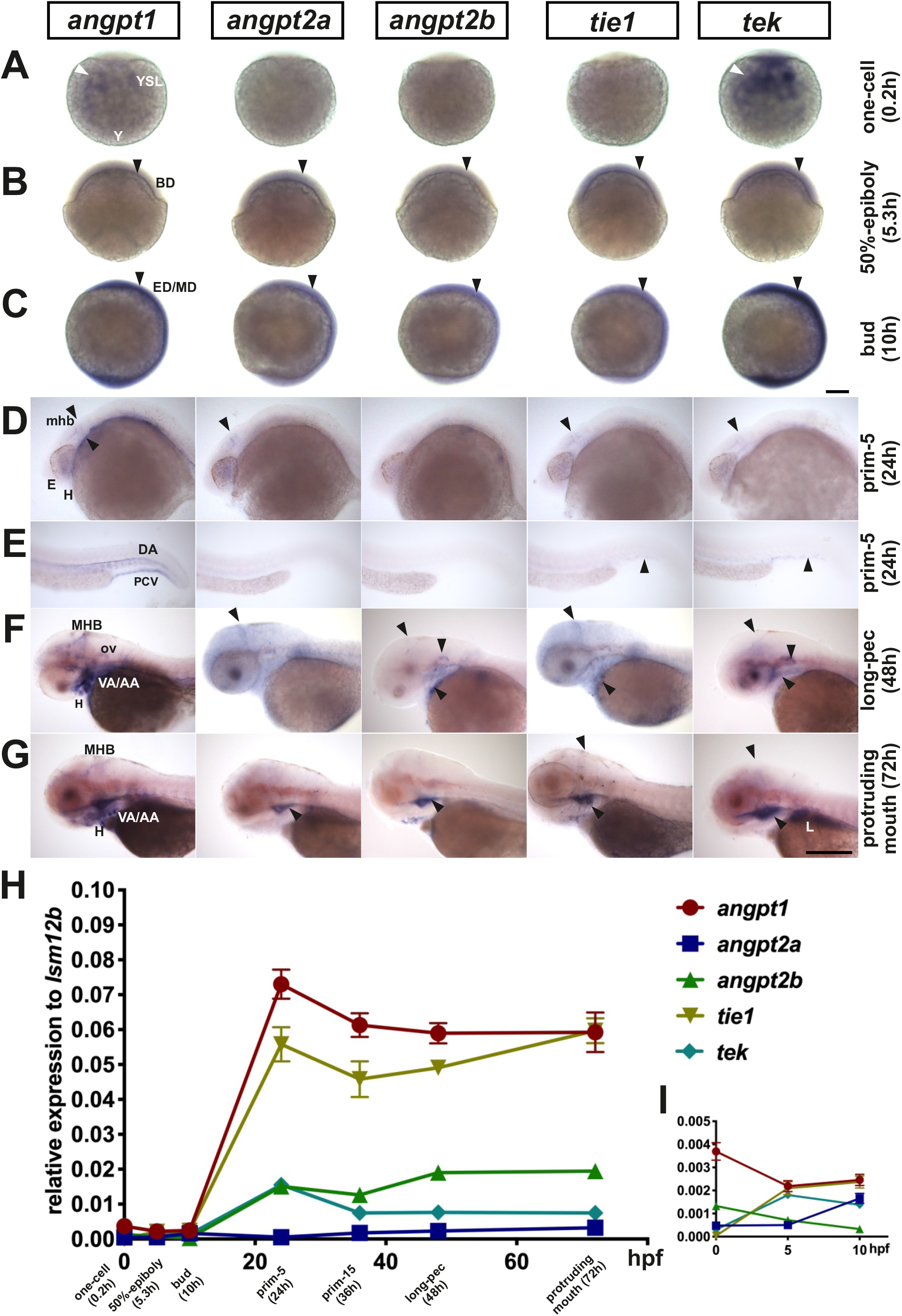
Spatiotemporal expression of zebrafish angiogenic factors, *angpt1, angpt2a, angpt2b, tie1 and tek*. Expression patterns of angiogenic factors from 7 developing stages (A) one-cell stage, (B) 50% epiboly, (C) bud, (D) head of prim-5, (E) trunk of prim-5, (F) long-pec and (G) protruding mouth (hatching period) by the whole-mount *in situ* hybridization. (H) Quantification of relative mRNA levels by qPCR through the one-cell stage to the hatching period. (I) mRNA levels during maternal-to-zygotic transition. Quantification of mRNA levels from the one-cell stage to the bud stage. Arrowheads indicate that angiogenic factors appear in blastoderm (BD), ectoderm/mesoderm (ED/MD), yolk and yolk syncytial layer (YSL). From pharyngula to hatching period, arrows display that signals appear in eyes (E), heart (H), liver (L), midbrain-hindbrain boundary (MHB), dorsal aorta (DA), branchial arch (AA) and posterior cardinal vein (PCV). Data are shown in mean ± SD (N=3 groups per stage, 20-30 embryos in one group) by one-way ANOVA with post-hoc Tukey’s test. Scale bar is 200 μm.

### C to T mutation in the coiled-coil domain of angpt1 causes loss of angpt1 function

To investigate the biological functions of angpt1 during embryonic neurogenesis, we carried out loss-of-function experiments using the heterozygous *angpt1* mutant line generated by TILLING (Targeting Induced Local Lesions IN Genomes) method. The *angpt1*^*sa14246*^ allele contains a C>T substitution. It introduced a premature stop codon at Q261, resulting in a truncated protein missing the entire fibrinogen-C-terminal domain, the region in which *Angpt1* binds to the *Tek* receptor (Fig. 2A). The mutation region was cloned, and the mutated nucleotide was confirmed by sequencing (Fig. 2B). Additionally, the genotype was determined at the earliest at 2-3 dpf by tail clipping using the High-Resolution melting curve analysis (HRM) (Fig. 2C), which was routinely used to genotype embryos and adult fish in this study. The genotypes from the same clutch followed the Mendelian distribution with approximately 25% inherited wild-type alleles (WT, *angpt1*^*+/+*^), 50% of heterozygous alleles (HET, *angpt1*^*+/−*^), and 25% of recessive mutants (KO, *angpt1*^*−/−*^).

**Fig. 2.**
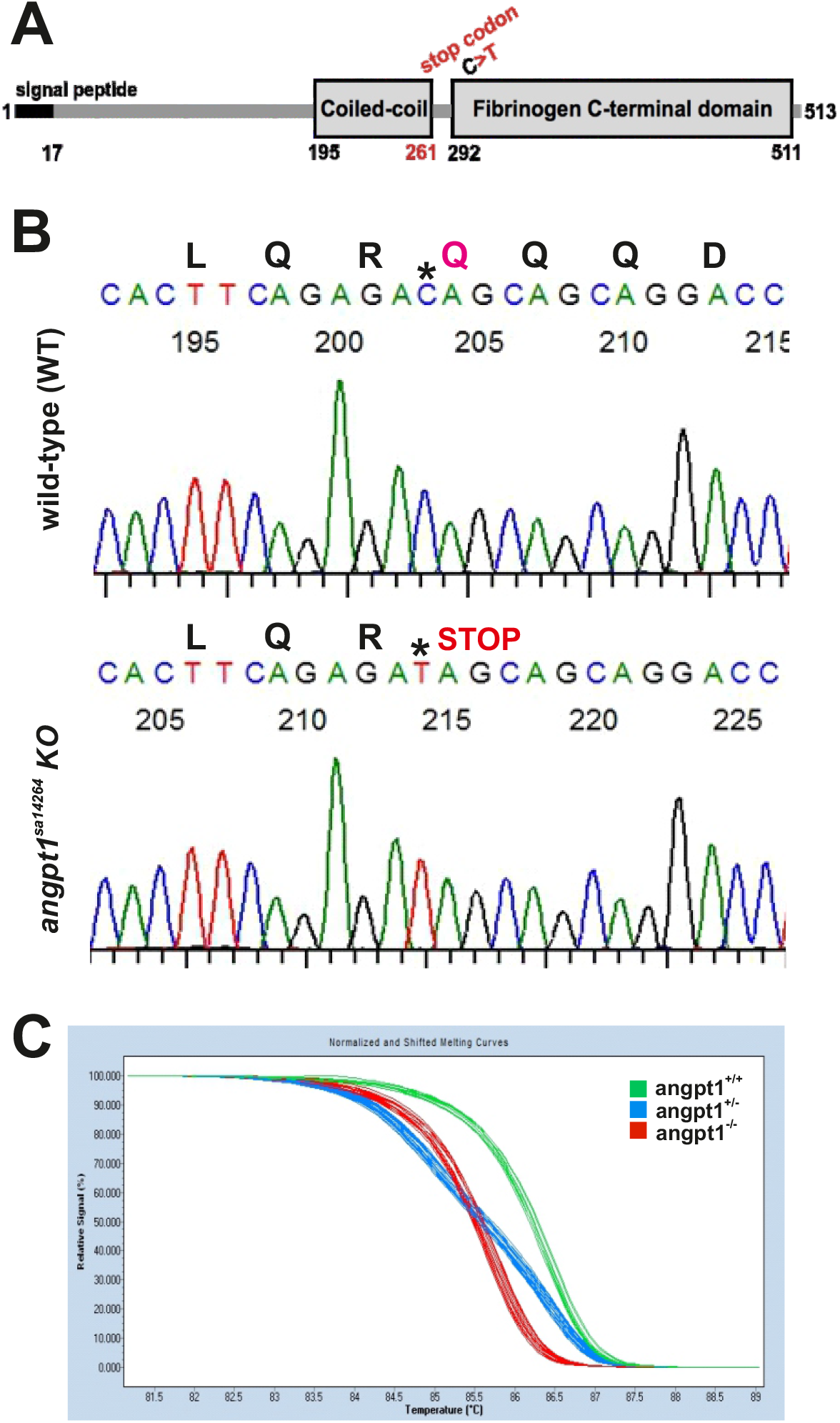
C to T mutation in the coiled-coil domain of *angpt1* gene causing a premature stop codon. (A) A scheme of angpt1 protein domains containing a signal peptide, coiled-coil and fibrinogen C-terminal domains. (B) The wild-type sibling and *angpt1*^*sa14266*^ mutant sequence chromatograms show that the *angpt1*^*sa14264*^ KO allele carries a nonsense mutation resulting in Q261 to a stop codon (*) in the coiled-coil domain. (C) The genotype of 3-dpf tail clips of *angpt1*^*sa14264*^ embryos generated by inbreeding heterozygous *angpt1*^*+/sa14264*^ parents. The high-resolution melting curve analysis distinguishes wild-type *angpt1*^*+/+*^ (WT), heterozygous *angpt1*^*+/−*^ (HET) from homozygous *angpt1*^*−/−*^ (KO) embryos.

### Embryonic lethality of loss of functional angpt1

At 36 hpf, one distinguishable defect was found in the clutch of the individual embryos produced by *angpt1*^*+/−*^ parents (Fig. 3A). *angpt1*^*−/−*^ embryos showed slower circulation with blood cells accumulated in the caudal vein. From 48 hpf outwards, approximately 25% of embryos showed pericardial edema (Fig. 3B). At 48 hpf, a significantly slower heartbeat rate than in control larvae (Fig. 3C), a slower blood flow velocity (Fig. 3D), and an impaired blood circulation in the caudal artery and veins (Fig. 3E) was found in *angpt1*^*−/−*^ embryos. At 72 hpf, *angpt1*^*−/−*^ embryos displayed more severe cardiovascular circulation impairments and pericardial edema. The smaller size of body (length) (Fig. 3F), brain (Fig. 3G), and eyes (Fig. 3H) were observed in homozygous mutant embryos. These phenotypic defects became progressively more severe after this time point, and *angpt1*^*−/−*^ embryos died at 5-6 dpf.

**Fig. 3.**
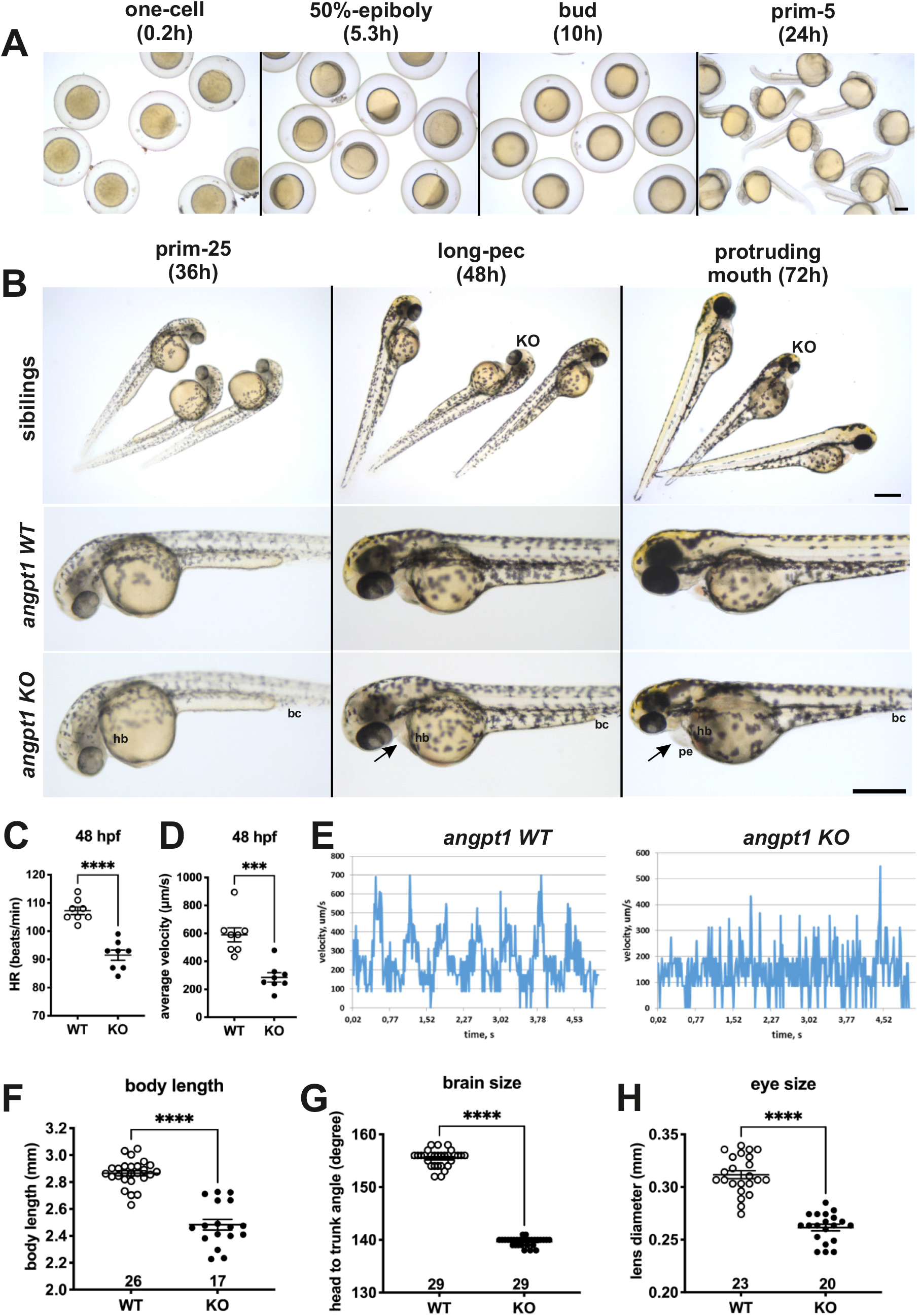
Aberrant embryonic brain and cardiovascular development of *angpt1* KO embryos at 48 hpf. All *angpt1*^*sa14264*^ samples older than 3 dpf were genotyped. 36-hpf and 48-hpf *angpt1*^*−/−*^ larvae (*angpt1 KO*) were defined primary by its distinctly failing cardiovascular circulation and then confirmed by genotyping. The 36-hpf and 48-hpf *angpt1* WT embryos were defined by the regular phenotype without genotyping. (A) Brightfield images show the normal appearance of *angpt1*^*−/−*^ (angpt1 KO) and its wild-type like siblings before 48 hpf. (B) Brightfield images show the *angpt1* KO embryos display cardiac edema at 72 hpf. The 3-dpf *angpt1* KO embryos show severe cardiac edema and smaller size of eye, head and body length compared with the wild-type like siblings. (C) Quantification of heartbeats in min (hb) and (D) the average velocity of blood circulation (bc) at 48 hpf. (E) Velocity profiles of the blood cells circulating in the dorsal aorta of *angpt1* WT and KO embryos at 48hpf. (F) Quantification of body length, (G) brain size, and (H) eye size of the *angpt1* WT siblings and *angpt1 KO* embryos at 3 dpf. Arrow indicates pericardial edema (pe). Sample numbers are sshown in the graphs. Data are shown in mean ± SEM. *p<0.05 and ***p<0.001 by Student’s t-test. Scale bar is 200 μm.

### Angpt1 deficiency causes anemia and aberrant patterning of cerebrovascular development

To study the hematopoietic development in the *angpt1*^*−/−*^ mutants, 3-dpf embryos were stained with o-dianisidine to detect hemoglobin containing cells. In the *angpt1*^*−/−*^ mutants, an intense hemoglobin staining was seen in the caudal artery and veins. In contrast, a robust reduction of o-dianisidine stainable cells was found in the heart, primary head sinus, and ventral aorta (Fig. 4A) compared with WT siblings. Moreover, brain haemorrhage was observed in 30% (21/68) of the 3-dpf *angpt1*^*−/−*^ mutants (Fig. 4A). To analyze the effects of loss of *angpt1* on the formation of the brain vasculature, confocal live images of microangiography were obtained from 3-dpf *angpt1*^*−/−*^ mutants and *angpt1* WT sibling. Remarkably, the *angpt1*^*−/−*^ larvae displayed severe defects mainly in the middle mesencephalic central artery and primordial hindbrain channel (Fig. 4B), but no overt morphological impairments were detected in the dorsal aorta, intersegmental vessels, caudal artery/veins in *angpt1* mutants despite the accumulation of erythrocytes in the caudal vein in the *angpt1*^*−/−*^ mutants (Fig. 4C), suggesting that *angpt1* plays a more crucial role in the regulation of the sprouting pattern of cerebrovascular development than in the trunk vessels during embryogenesis.

**Fig. 4.**
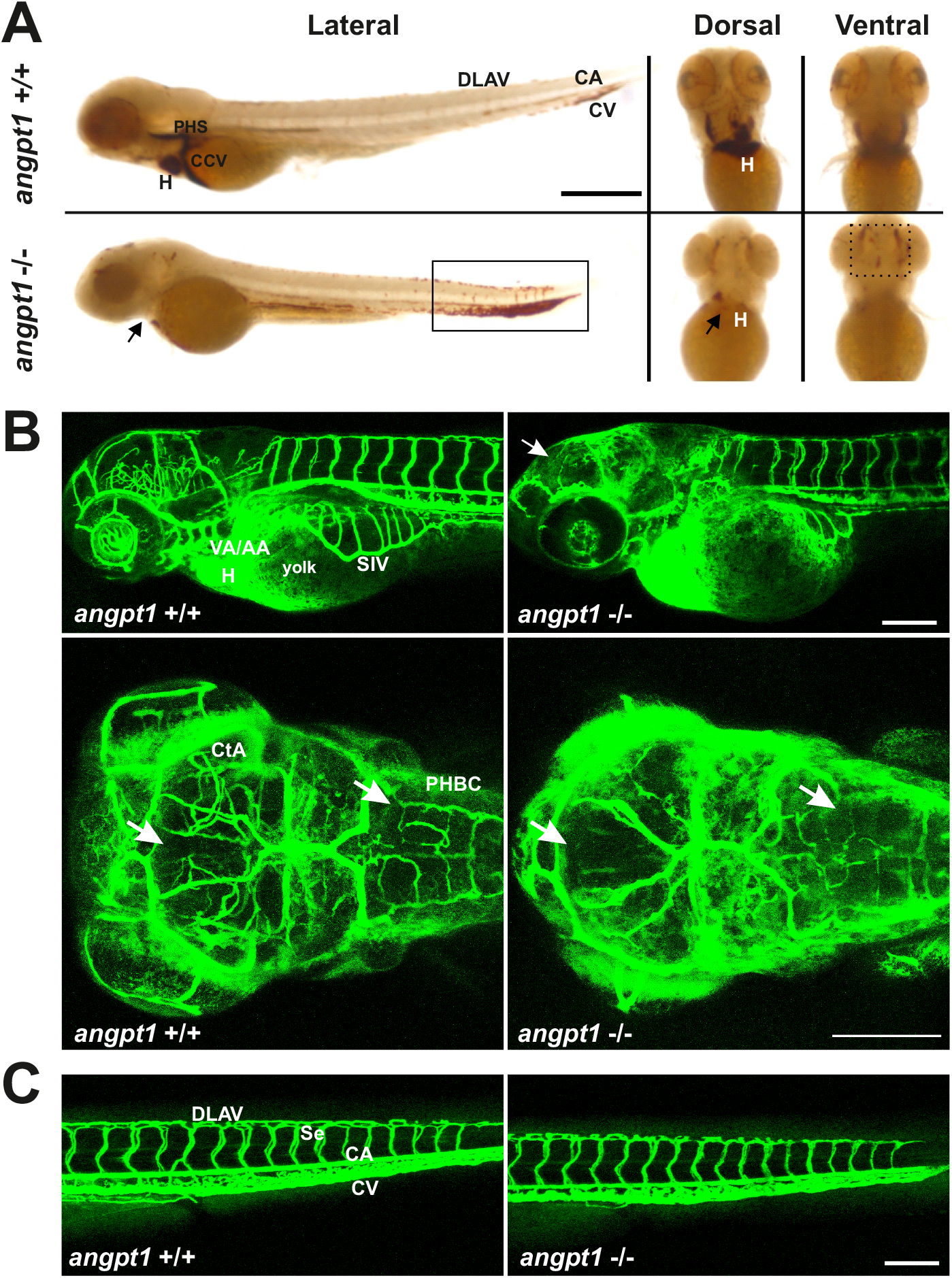
Brain haemorrhage and impairments of cerebrovascular development in *angpt1* KO embryos. (A) The blood phenotype of 3-dpf *angpt1* KO embryos shown by whole mount o-dianisidine staining. Lateral views and ventral views show that *angpt1* mutants lack hemoglobinized erythrocytes in the heart (arrow) and have impaired blood circulation with the accumulation of blood cells in the aorta and ventral tail (rectangle) and brain haemorrhage (dash line rectangle). (B) Microangiography of the vascular formation in *angpt1*^*−/−*^ mutants and WT siblings. Microangiography was done by injecting fluorescent dye directly into the sinus venosus in 3-dpf genotyped embryos. Lateral views and dorsal views of maximal intensity projections of confocal z-stacks images indicate that loss of *angpt1* causes the malformation of forebrain and hindbrain vascular patterning. (C) No apparent defects in the trunk vasculature of *angpt1* mutants. CA, caudal artery; CtA, central artery; CV, caudal vein; DLAV, dorsal longitudinal anastomotic vessels; H, heart; MCeV, middle cerebral vein; PHBC, primordial hindbrain channel; Se, intersegmental vessel; SIV, subintestinal vein; VA/AA, ventral aorta/ branchial artery. CA, caudal artery; CCV, common cardinal vein; CV, caudal vein; DLAV, dorsal longitudinal anastomotic vessel; H, heart; PHS, primary head sinus. Scale bar is 200 μm.

### Defective neurogenesis in 36-dpf *angpt1* mutant larvae

Apart from a vital function in angiogenesis, our previous study showed that angpt1 was closely associated with the regulation of embryonic neurogenesis, such as regulating the brain size through the notch1 signaling in the teleost fish (Chen et al., 2015). To further investigate the effect of loss of *angpt1* on early embryonic neurogenesis, we first studied the expression of relevant marker genes by WISH in 36-hpf *angpt1* KO and *angpt1* WT embryos. Compared with its *angpt1* WT siblings, the *angpt1* KO embryos showed a dramatically reduced *angpt1* expression in the head, in the caudal perivascular artery and veins (Fig. 5A), confirming that *angpt1* mRNA containing premature translation termination codon was degraded via the nonsense-mediated mRNA decay pathway. However, faint signaling was still detected in the 36-hpf *angpt1* KO embryos, possibly due to residual expression of maternal *angpt1* or aberrant mRNA partially hybridized with the riboprobe because 3-dpf *angpt1*^*−/−*^ embryos showed no *angpt1* signals (Fig. 6A). *tek* expression showed only modest differences in the head and the caudal perivascular region between *angpt1* WT and *angpt1* KO siblings at this stage (Fig. 5B). notch1a expression showed no dramatic difference in the *angpt1* KO and WT embryos (Fig. 5C). Notably, *nestin* (a marker for neuronal precursors in the developing central nervous system (CNS) giving rise to both neurons and glia) and *wnt1* (regulating midbrain and hindbrain formation) mRNA, were downregulated particularly in the midbrain-hindbrain boundary and in the rhombomeres in the *angpt1* KO mutants (Fig. 5D and 5E), suggesting that angpt1 has a significant impact on the early brain morphogenesis in the zebrafish.

**Fig. 5.**
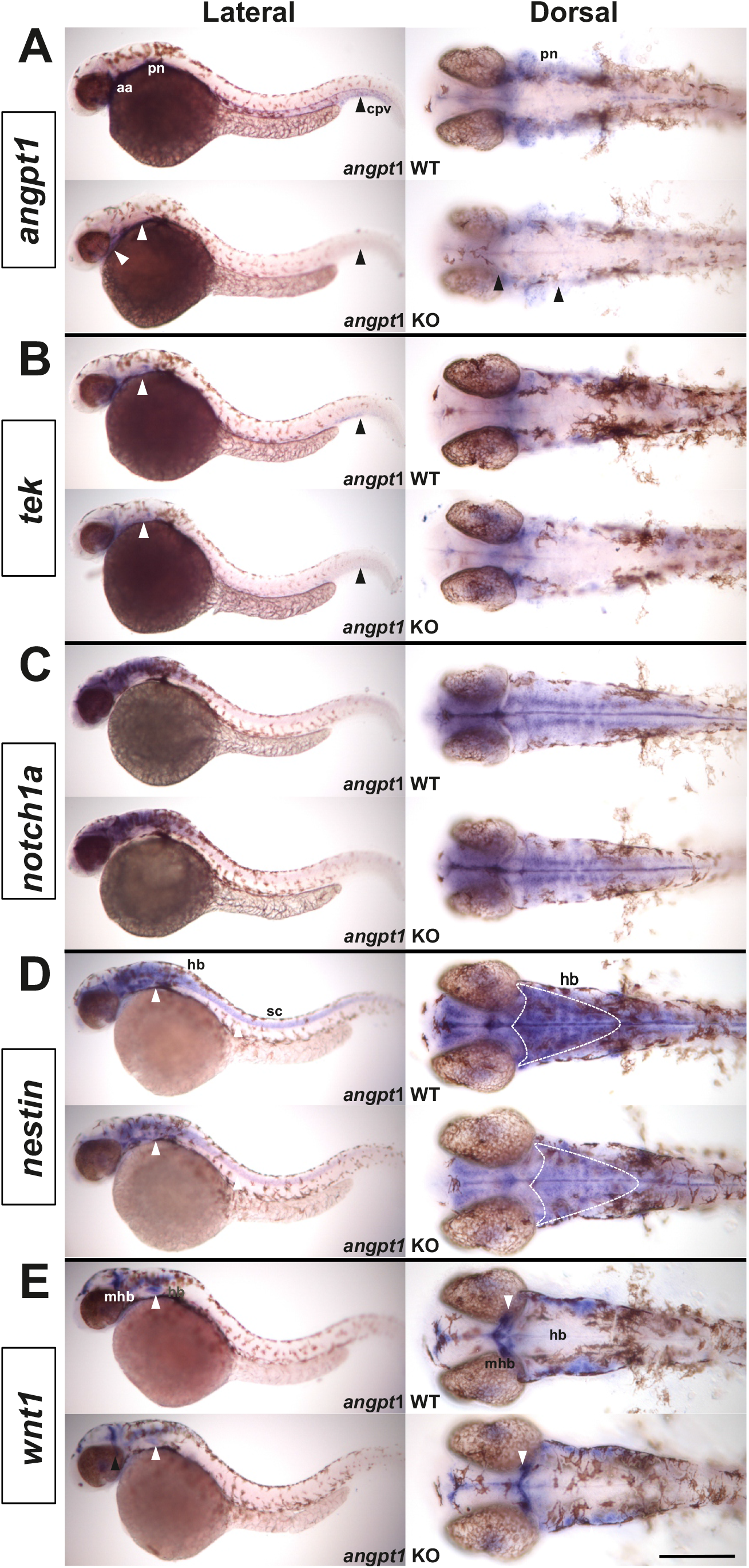
Spatial mRNA distribution of genes involved in angiogenesis and patterning processes in 36-hpf *angpt1* mutants. (A) *angpt1* and (B) *tek* expression are reduced in aortic arches (aa), and caudal perivascular region (cpv) in the *angpt1* KO group. (C) *notch1a* expression showed no apparent difference between the *angpt1* KO and its sibling WT group. (D) *nestin* expression is lower in *angpt1*^−/−^ embryonal hindbrain (hb) and spinal cord (sc) than in the *angpt1*^+/+^ embryos. (E) *wnt1* mRNA signal was lower in the midbrain-and-hindbrain boundary (mhb) and the hindbrain in *angpt1*^−/−^ than in *angpt1*^+/+^ embryos. Arrowheads indicate the signal difference between the WT embryos and the *angpt1* KO group. N=5 in each genotyped group. Scale bar is 200 μm.

**Fig. 6.**
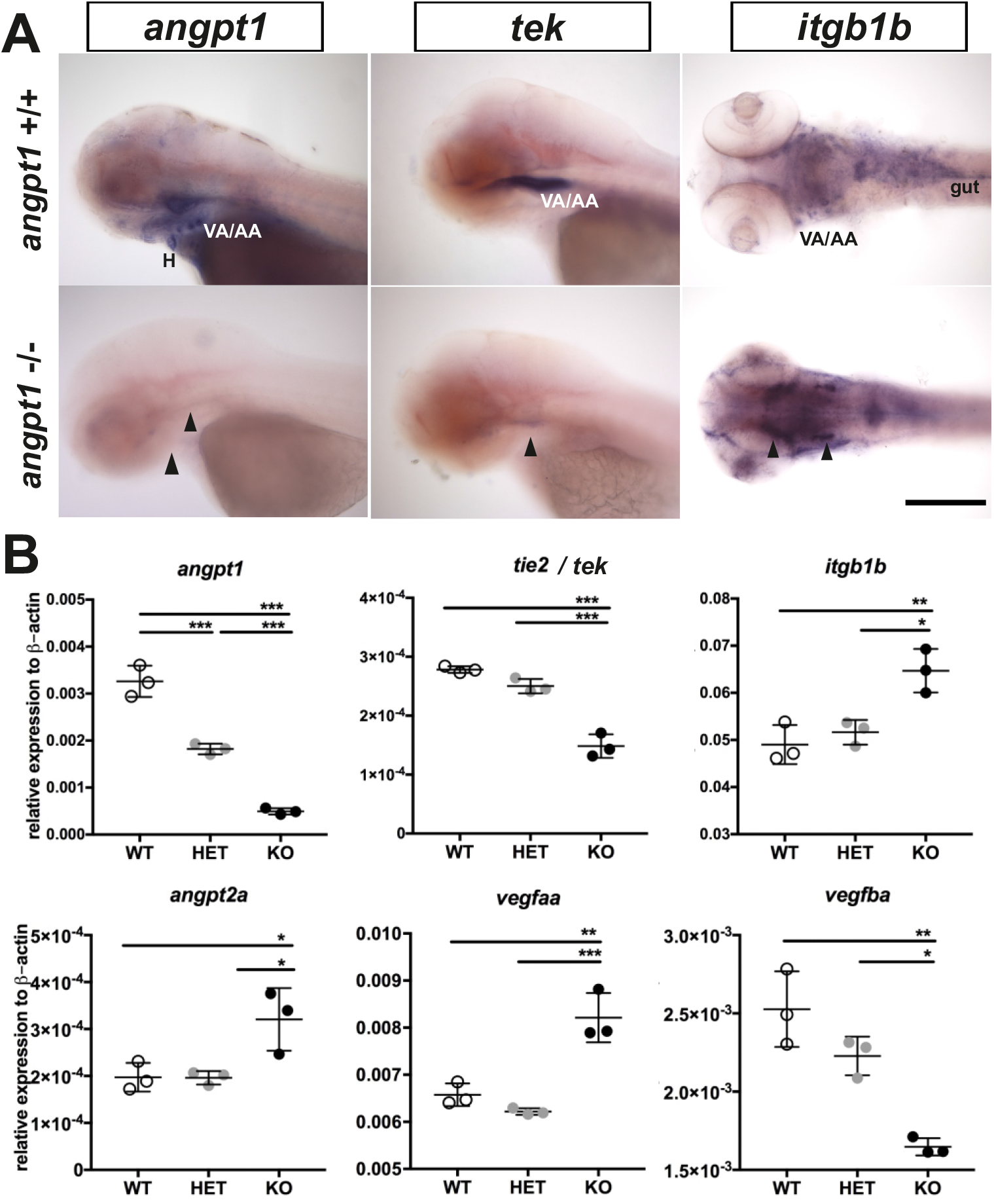
Expression of *angpt1*, *tek* and *itgb1b* mRNA in 3-dpf *angpt1* mutants. (A) The mRNA distribution of *angpt1* and *tek* is nearly undetectable in *angpt1* KO mutant, whereas expression of *itgb1b* mRNA is significantly higher in *angpt1* mutants than WT siblings. (B) Quantification of mRNA levels of *angpt1, tek, itgb1b, angpt2a, vegfaa and vegfba* by qPCR. Each embryo was genotyped with HRM analysis. N=6 per group for WISH. N =3 per group and 10-pooled embryos in one group for qPCR. H, heart; VA/AA, ventral aorta/ branchial arch. Arrowheads indicate sites of differential gene expression. *p<0.05, **p<0.01 and ***p<0.001 by Student’s t-test.

### Angiogenic factors in 3-dpf *angpt1* mutants

Using 3-dpf genotyped embryos, we further confirmed that the phenotypic abnormalities appearing in *angpt1* KO embryos accurately correlated with the genotyping outcome by WISH (Fig. 6A) and the qPCR (Fig. 6B). Compared with *angpt1*^*+/+*^ siblings, only 55.8 % and 15% of remaining *angpt1* mRNA were detected in the *angpt1*^*+/−*^ and *angpt1*^*−/−*^larvae (Fig. 6B), respectively, confirming that the truncated form of the *angpt1* transcript was targeted for the nonsense-mediated decay surveillance pathway (NMD). Concomitantly with the *angpt1* downregulation, a significant decrease of *tek* expression (specific binding receptor of angpt1) was found in the *angpt1*^*−/−*^ mutants (Fig. 6A and 6B). In contrast, *itgb1b* (the integrin family as a potential critical receptor of angpt1) and *angpt2a* (a context-dependent antagonist of *angpt1*) were significantly upregulated in the *angpt1*^*−/−*^ larvae. Intriguingly, the expression level of *vegfaa* and *vegfba*, essential factors for vascular remodeling, were upregulated and downregulated, respectively, in the *angpt1*^*−/−*^ larvae. These dynamic alterations of angiogenic factor expression patterns may represent compensating efforts to cover the loss of functional *angpt1* commending to stabilize neurovascular formation and angiogenesis during embryonic development.

### Markers of neurogenesis in *angpt1*^*sa14264*^ embryos

Proper coordination of neurogenesis and angiogenesis is crucial for the normal construction of the CNS and neurovascular system during embryonic development (Ward and Lamanna, 2004). We further investigated relevant marker genes for embryonic neurogenesis using 3-dpf genotyped sibling larvae by WISH (Fig. 7A) and qPCR (Fig. 7B). The early neural marker *notch1a* and *nestin* expression were reduced in the head, branchial arches, and hindbrain of the angpt1^−/−−^larvae compared with their *angpt1*^*+/+*^ siblings (Fig.7A). The mRNA levels of *notch1a* and *nestin* were significantly downregulated in the *angpt1*^*−/−*^ larvae compared with *angpt1*^*+/+*^ and *angpt1*^*+/−*^ siblings (Fig. 7B). The expression level of *pcna* (proliferating cell nuclear antigen, polymerase delta auxiliary protein), *wnt1* and *wnt10b* (Table2; wnt family, regulators of cell fate and patterning during embryogenesis), and *SRY-box containing gene 2* (*sox2*) were expressed in significantly lower levels in the *angpt1*^*−/−*^ larvae than in *angpt1*^*+/+*^ and *angpt1*^*+/−*^ siblings. In contrast, *wnt2bb* expression was upregulated in the *angpt1*^*−/−*^ larvae (Fig. 7B). To find out whether specific neuron populations are affected in *angpt1*^*−/−*^ larvae, markers of aminergic neurons were studied. Interestingly, expression of *tyrosine hydroxylase 1* (*th1*, a marker of dopaminergic and noradrenergic neurons) was significantly downregulated, while the expression levels of *th2* (a marker of non-overlapping dopaminergic neurons) and *histidine decarboxylase* (*hdc*, a marker of histaminergic neurons) were unaltered in the *angpt1*^*−/−*^larvae (Fig. 7C). The mRNA expression levels of glial markers (*gfap* and *apoEb*) were significantly increased in the *angpt1*^*−/−*^ mutants (Fig. 7D), and the number of *apoeb*-positive cells was significantly higher in the midbrain of *angpt1*^*−/−*^ larvae than in WT siblings (Fig.7A, 7E). Of neurogenic markers, genes representing proliferating and neural progenitor cells were downregulated in the 36-hpf and 3-dpf *angpt1*^*−/−*^ embryos (Fig 5 and 7), suggesting that loss of functional *angpt1* has a significant effect on the early neurogenesis.

**Table 2.**
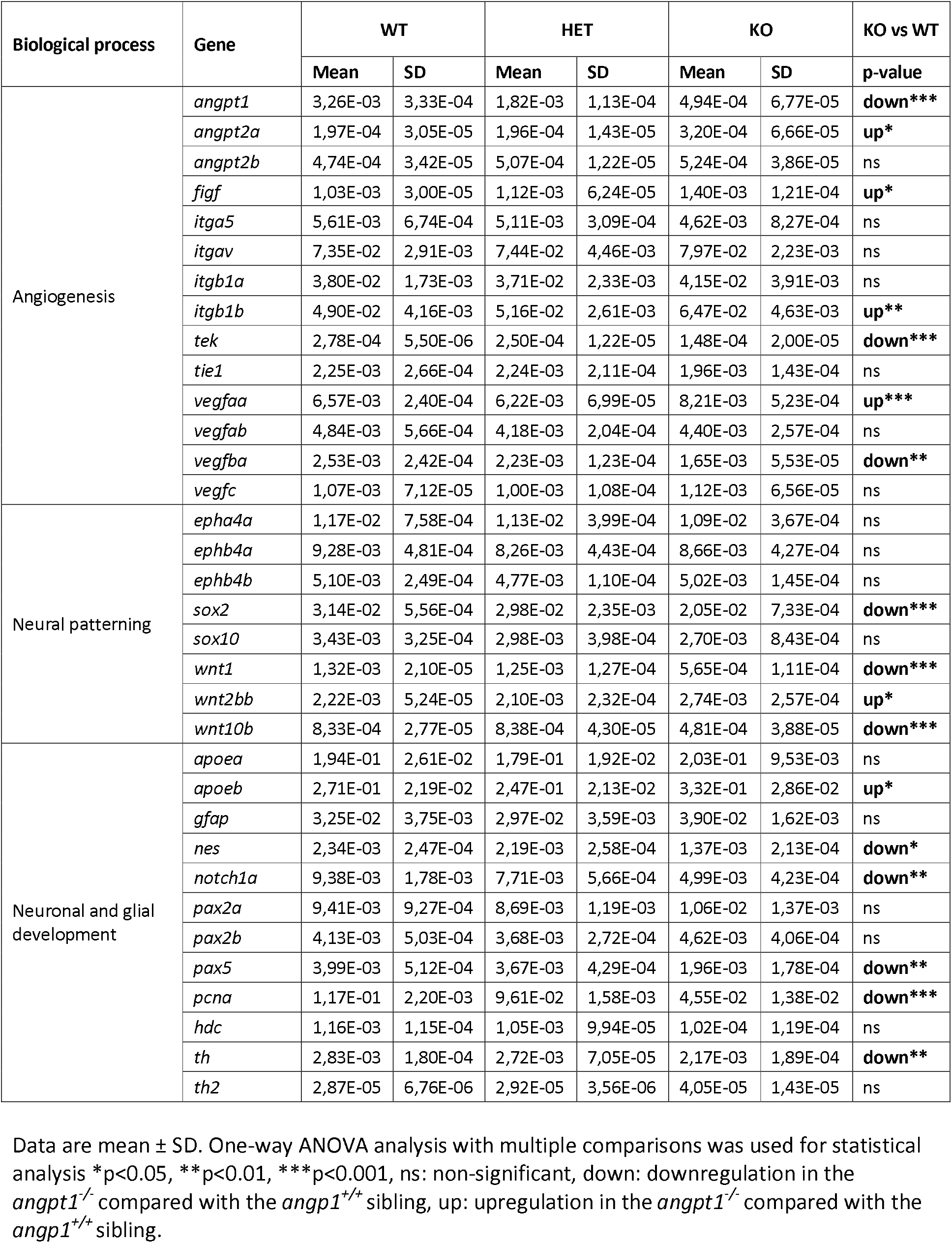
Summary of RT-qPCR results of 3-dpf *angpt1*^*−/−*^ embryos.

**Fig. 7.**
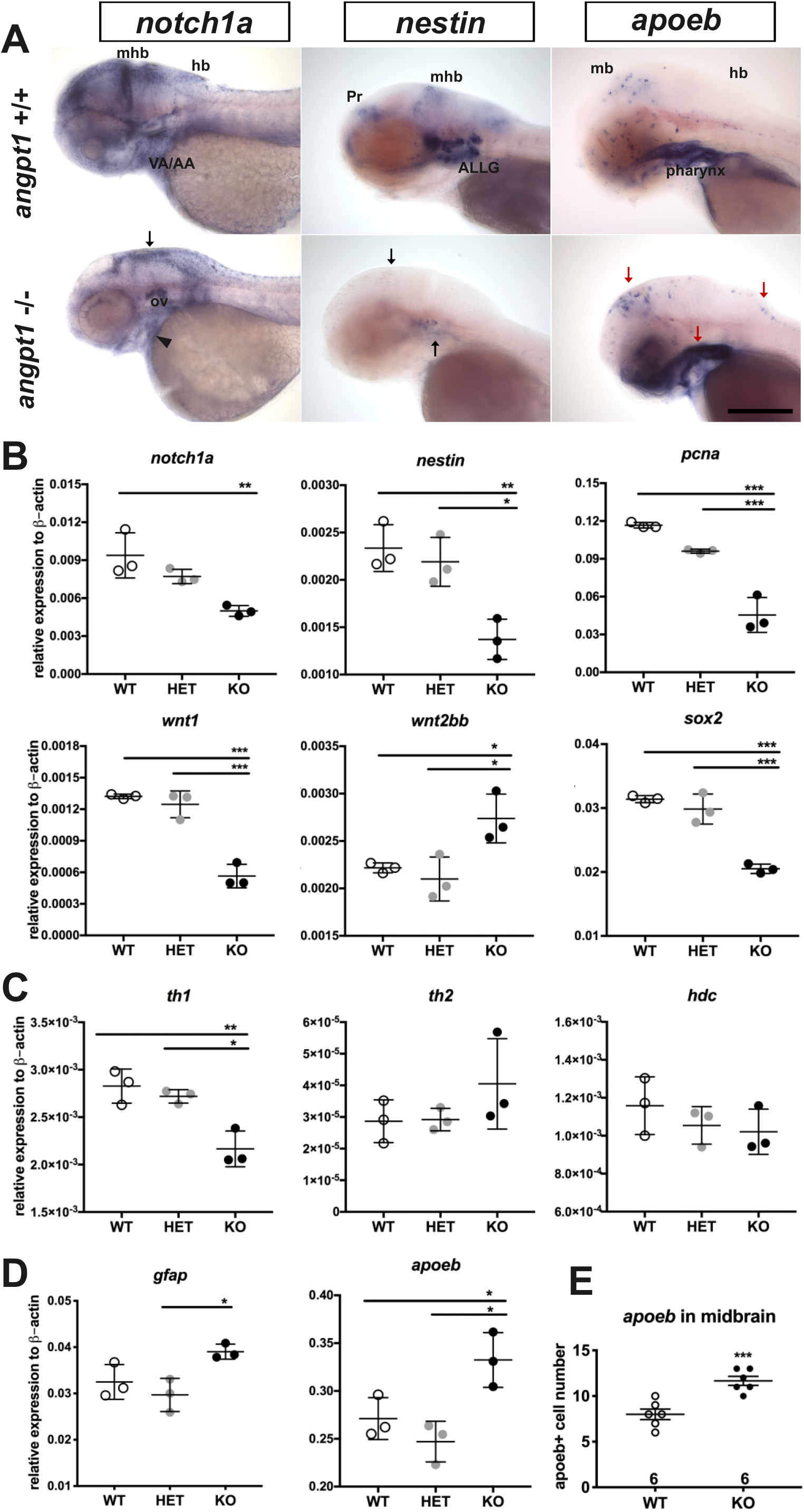
Relative expression levels of neurogenesis markers in 3-dpf *angpt1* mutants. (A) *notch1a* and *nestin* expression is lower, whereas *apoeb* is significantly higher in the midbrain brain (mb), hindbrain (hb) and ventral aorta/ branchial arch (VA/AA) in the *angpt1* ^*−/−*^ than in *angpt1*^*+/+*^ embryos by WISH (n=5 in each group) (B) Quantification of mRNA expression levels of genes involved in embryonic neurogenesis in 3-dpf embryos. The mRNA expression of *notch1a, nestin, pcna, wnt1, and sox2a* is lower, but the *wnt2bb* is higher in the *angpt1* mutants than in WT embryos. (C) *th1* mRNA expression is lower in *angpt1*^−/−^ than in *angpt1*^*+/+*^ embryos, but *th2* and *hdc* mRNA expression is not different between the genotypes. (D) *gfap* and *apoeb* expression is higher in the *angpt1* KO larvae compared with WT siblings. (E) Quantification of *apoeb*-positive cell numbers in the midbrain. Arrows indicate the sites of differential gene expression between WT and *angpt1* KO. N=6 per group for WISH. N=3 per group and 10-pooled embryos in one group for qPCR. ALLG, anterior lateral line ganglion; Pr, pretectum; The data represent mean ± SD of three independent experiments (n=3 pre-group and 10-pooled embryos in one group). *p<0.05, **p<0.01 and ***p<0.001 by Student’s t test.

### Downregulation of proliferation and upregulation of astrogliogenesis/gliagenesis in *angpt1* and *itgb1b* mutant brain

Unlike the gross cardiovascular phenotype appearing in the *angpt1*^*sa14264*^ mutant and reported *itgb1b*^*mi371*^ mutant (Iida et al., 2018), the *tek*^*hu1667*^ mutant fish grow naturally without overt cardiovascular defects (Gjini et al., 2011; Jiang et al., 2020). It has been reported that besides Tek, Angpt1 can bind to integrins and activate similar pathways as the angpt1-tek does (Chen et al., 2009a), which renders it possible that the angpt1-itgb1b signaling pathway may play a more vital role than the recognized angpt1-tek pathway in the regulation of embryogenesis. To provide detailed evidence that angpt1 and itgb1b are involved in the regulation of proliferation during neurogenesis, we conducted the saturation-labeling EdU incorporated analysis followed by staining with the glial marker GFAP on 3-dpf *angpt1* and *itgb1b* mutant larvae. A significant decrease of EdU-positive cells was found in the proliferative zones in both *angpt1* (Fig. 8A) and *itgb1b* KO larvae (Fig. 8B). In contrast to the diminished proliferation, a robust increase of GFAP-positive signals appeared in the forebrain and hindbrain of *angpt1*^*−/−*^ and *itgb1b*^*−/−*^ larvae compared with their WT siblings, indicating that angpt1 and itgb1b have a vital role in the regulation of proliferation during embryonic neurogenesis.

**Fig. 8.**
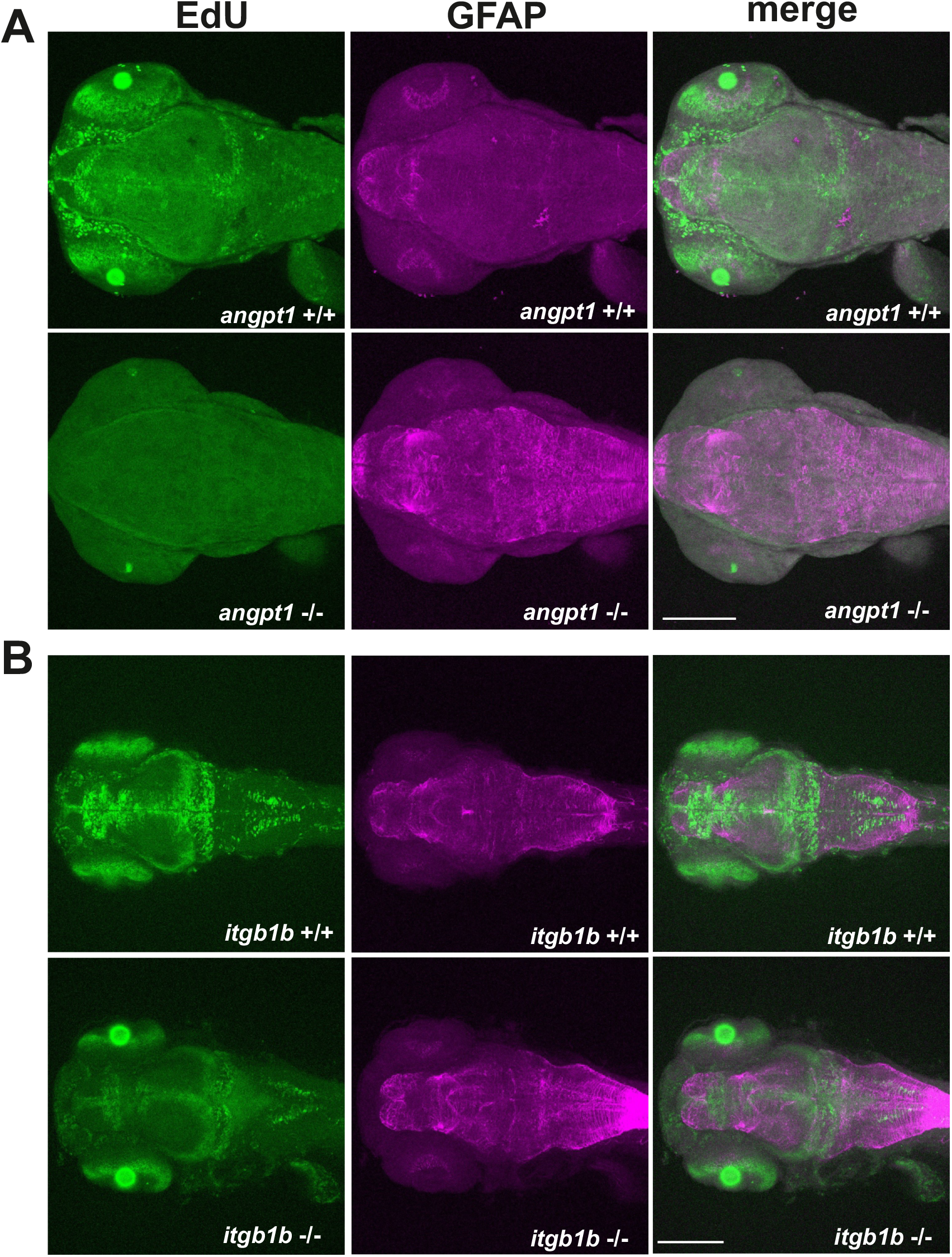
Downregulation of proliferation but upregulation of astrogenesis in *angpt1* and *itgb1b* mutants. Maximum intensity projections of confocal z-stack images of immunostaining of GFAP following the EdU proliferation assay in (A) 3-dpf *angpt1*^*+/+*^ and *angpt1*^−/−^ embryos and (B) 3-dpf *itgb1b*^+/+^ and *itgb1b*^−/−^ embryos. Proliferating cells indicated in green, and GFAP-positive signals in magenta. N=5 per group. Scale bar is 200 μm.

### Distortion of *krox20* patterns in the hindbrain of *angpt1* and *itgb1b* mutants

Wnt signaling pathways play a crucial role in the regulation of rhombomere segmentation during embryogenesis (Riley et al., 2004). Wnt1 is mainly involved in the formation of the midbrain-hindbrain boundary and hindbrain patterning (Duncan et al., 2015). As mentioned above, in the *angpt1*^*−/−*^ mutants, expression levels of *wnt1, wnt2bb*, and *wnt10b* were significantly altered (Table 2), and these substantial alterations may cause impairments of hindbrain patterning. We further found that *krox20*/ *egr2b*-positive cells (encoding a transcription factor playing a pivotal role in hindbrain segmentation) in the hindbrain and lateral line nerves were mostly missing in the *angpt1*^*−/−*^ mutants (Fig. 9A); similarly, fewer and disorganized *egr2b*/*krox20*-positive signals were found in the *itgb1b*^−/−^ larval hindbrain compared with WT siblings (Fig. 9B).

**Fig. 9.**
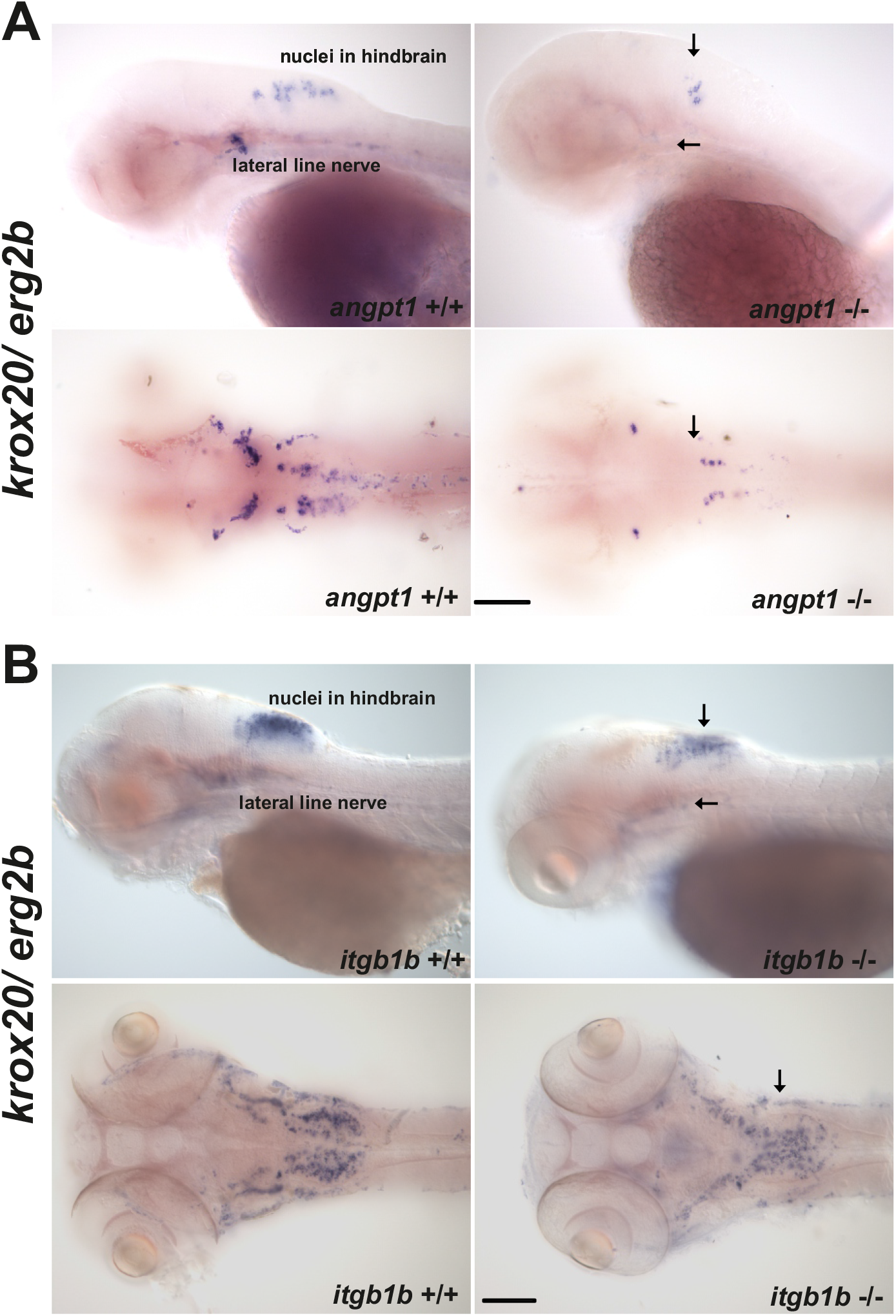
Abnormal expression of *Krox20* in *angpt1* and *itgb1b* KO hindbrain. Expression of *krox20* mRNA by WISH in *angpt1* and *itgb1b* KO larval hindbrain (A) abnormal restricted expression patterns of *krox20* shown in the hindbrain in 3dpf *angpt1* mutants. (B) abnormally low expression of *krox20* in 4-dpf *itgb1b* mutants. N=5 each group. Arrows indicated the *krox20* -positive cells in the hindbrain and lateral line nerve. Scale bar is 200 μm.

### Impairments of retrograde labeling of reticulospinal neurons in *angpt1* and *itgb1b* mutants

The development of the hindbrain segmented into rhombomeres plays a crucial role in orchestrating the stereotypical organization of reticulospinal, somatomotor, and branchiomotor neurons. To further investigate whether the reticulospinal neurons were affected in *angpt1* and *itgb1b* mutant hindbrain, we performed retrograde labeling of neurons using the green fluorescent dye conjugated high-molecular-weight dextran (10,000Mw). Retrograde labeling showed a severe deficiency of reticulospinal neurons in the *angpt1*^*−/−*^ and *itgb1b*^*−/−*^ mutant hindbrain (Fig. 10D and 10E), which were in contrast to the normal patterning of their WT siblings (Fig.10A and 10B) and the *tek*^*−/−*^ mutant (Fig. 10C and 10F). On the other hand, some positive fluorescent cells were found in the *angpt1*^*−/−*^ and *itgb1b*^*−/−*^ mutant brains. This may be due to the breakdown of the cerebrovascular barriers and increased vascular permeability caused by loss of functional *angpt1* and *itgb1b*, or lacking active retrograde transports inside neurons in these mutants so that the labeling dye diffused into the cerebral ventricle system and labeled some cells ectopically. The breakdown of BBB also produced brain haemorrhage that appeared in both *angpt1*^*−/−*^ (Fig.4A) and *itgb1b*^*mi371*^mutants (Iida et al., 2018), but not in normally developed *tek* mutant fish brains (Gjini et al., 2011).

**Fig. 10.**
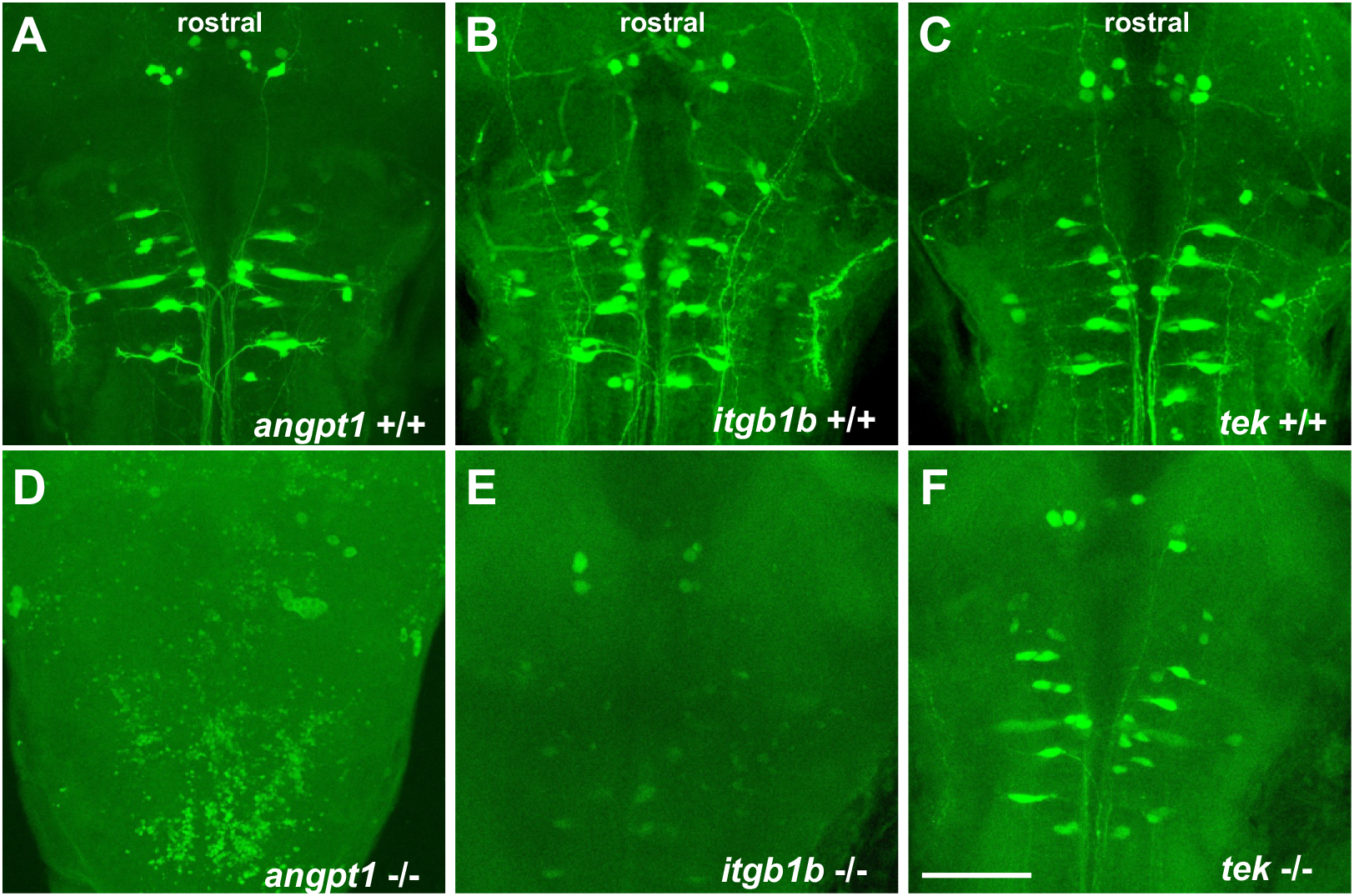
Impairment of reticulospinal neurons in *angpt1* and *itgb1b* mutants. Retrograde labeling of 4-dpf genotyped larvae reveals the deficiencies of reticulospinal neurons in *angpt1* KO and *itgb1b* KO brains. (A) *angpt1* WT (B) *itgb1b* WT (C) *tie2* WT (D) *angpt1* KO (E) *itgb1b* KO and (F) *tie2* KO. N=5 per group. Scale bar is 200 μm.

### Deficient dopaminergic and histaminergic neurons in *angpt1* and *itgb1b* mutant brains

In addition to angiogenic effects, angiogenic factors can function as neurotrophic factors, not only involved in neurogenesis but also protecting neuron loss against different types of insults. We then investigated some well-known neurotransmitter systems, dopaminergic and histaminergic populations in the 4-dpf larval brain. We quantified the cell numbers following immunostaining in the caudal hypothalamus, which contains the most representative neurotransmitter systems, including dopaminergic, GABAergic, histaminergic and serotonergic neuron groups. In the *angpt1*^*−/−*^ and *itgb1b*^*−/−*^ mutant brains, a significant decrease in the number of TH1-positive cells was evident in the caudal hypothalamus (Fig.11A, 11B). Likewise, a significant reduction of histamine-positive cells was found in both mutants (Fig. 11D, 11E). In contrast to these two mutants, the number of TH1-positive cells and histaminergic cells remained intact in *tek*^*−/−*^ mutants, similar to their WT siblings (Fig. 11C, 11F). The deficiency of dopaminergic and histaminergic populations in the *angpt1*^*sa14264*^ and *itgb1b*^*mi371*^ mutant brains indicates the importance of angpt1 and itgb1b for differentiation and maintenance of developing and mature neurons during development.

**Fig. 11.**
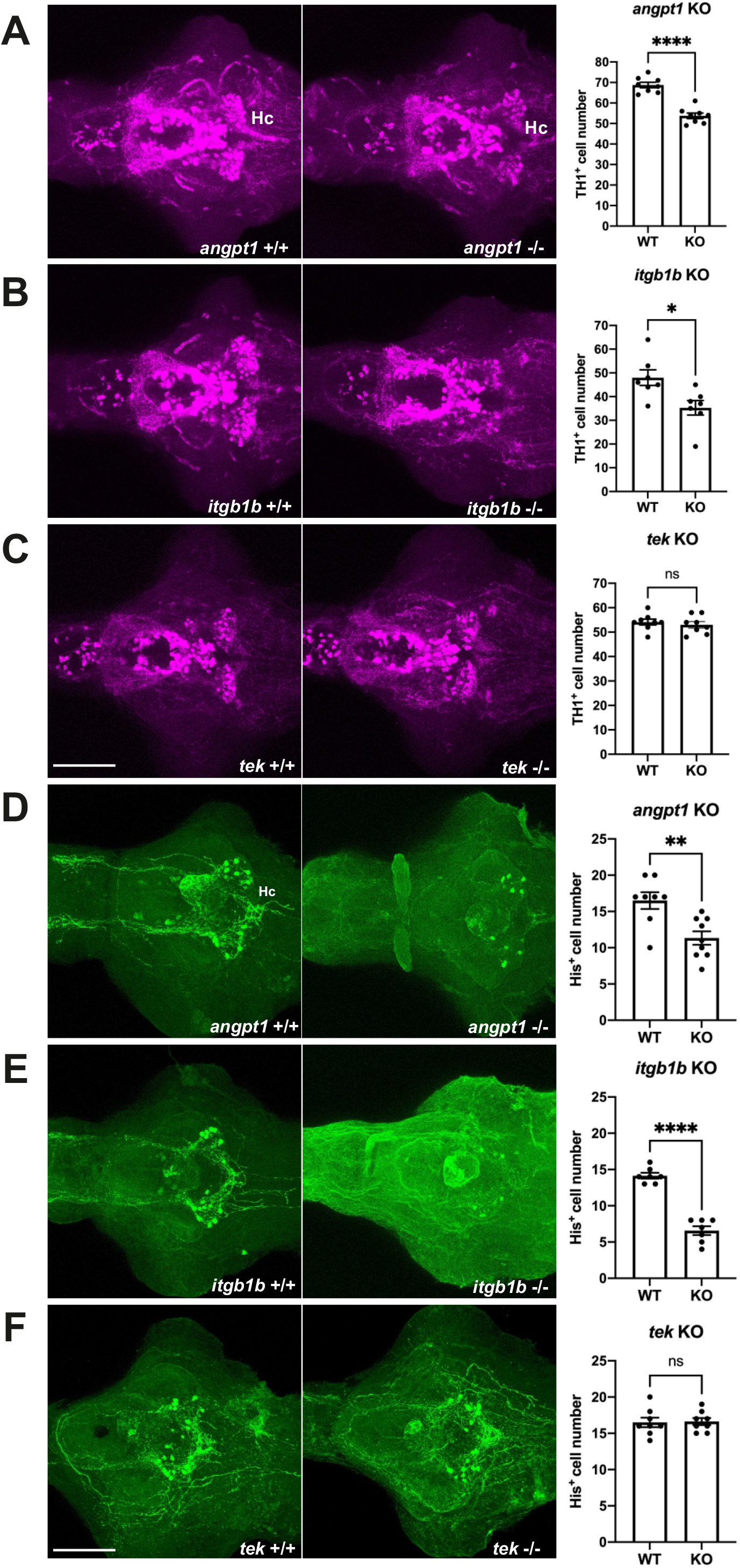
Deficient dopaminergic and histaminergic neurons in the hypothalamus of *angpt1* KO and *itgb1b* KO larvae. (TH1 immunostaining images of 4-dpf (A) *angpt1* KO, (B) *itgb1b* KO and (C) *tek* KO brains. Quantitative results show a significant decrease of TH1-positive cell number found in the caudal hypothalamus of *angpt1* KO and *itgb1b* KO brains. Histamine (His+) immunostaining images of 4-dpf (D) *angpt1* KO, (E) *itgb1b* KO and (F) tek KO brains suggests low cell numbers and deficient fiber networks. Quantification of His-positive cell numbers shows a decreased number in *angpt1* and *itgb1b* KO brains. Data represent the mean ± SEM. The student’s t-test was used for statistical analysis. *p< 0.05, **p<0.01 and ***p<0.001. Scale bar is 200 μm

### Overexpression of zebrafish *angpt1* promotes cardiovascular activity and neurogenesis

Loss of functional angpt1 revealed downside effects on embryonic neurogenesis. To study whether angpt1 alone is sufficient to regulate angiogenesis and neurogenesis during embryogenesis, we employed a gain-of-function system by injecting a transgene plasmid containing Tol2 transposase sites and encoded a full-length zebrafish *angpt1* driven by the ubiquitin promoter (*ubi*) (Mosimann et al., 2011) to overexpress *angpt1* ubiquitously *in vivo* along with a heart-specific marker, the cmlc2 promoter driving *gfp* expression. As a result, the transgene was randomly intergraded into the genome so that embryos with GFP expression in the heart, indicating successful transgene integration, were selected for further experiments. We showed with *angpt1* WISH that *angpt1* was ectopically overexpressed in the heart and trunk (Fig. 12A) although the body size (data not shown), brain length (data not shown), and average blood flow (Fig. 12D) were unaffected. The heartbeat rate (Fig. 12B) was higher in the *ubi*-driven *angpt1* fish than in the control-injected group. Furthermore, overexpression of *angpt1* led to ectopic vessel formation in the head and the trunk (Fig. 12C). An upregulation of *notch1a* was found in the brain ventricular zone in the *ubi*-driven *angpt1* group (Fig.12A). Moreover, quantification of mRNA levels confirmed that the *angpt1* mRNA level was 40-fold higher in the *ubi*-driven *angpt1* group than the control-injected one, and *tie1* expression was also upregulated concomitantly (Fig.12E). Notably, the *notch1a* expression was quantitatively upregulated as well. Concurrently, the expression of *gfap* was downregulated in the *ubi*-driven *angpt1* group (Fig. 12E). Overexpression of angpt1 revealed the anticipated opposite outcomes compared with the loss-of-functional *angpt1* effects shown in Fig. 7 and 8 in the *angpt1*^*−/−*^ fish, providing evidence that *angpt1* has a crucial role in the regulation of cardiovascular formation and neurogenesis signaling during embryogenesis.

**Fig. 12.**
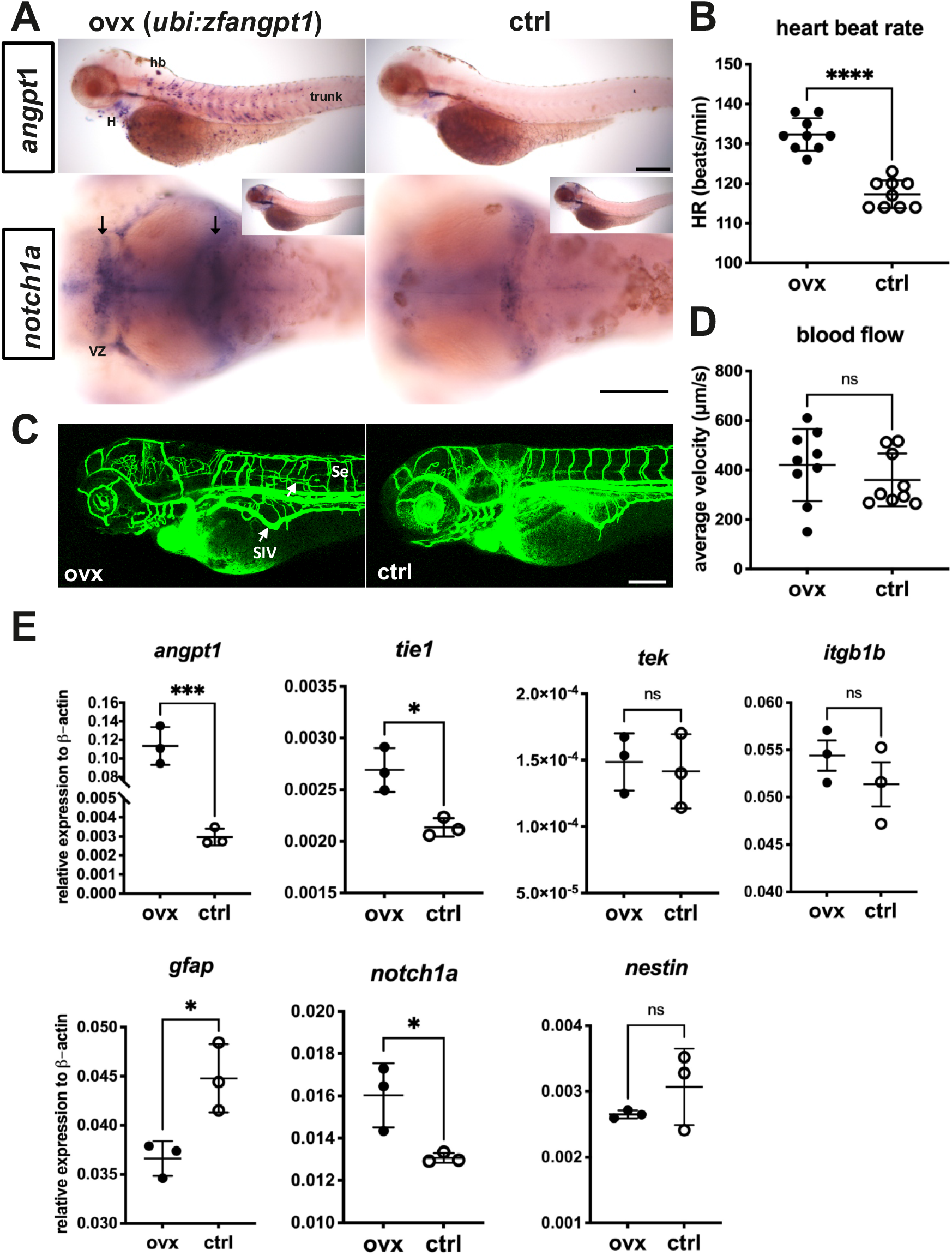
Ectopic expression of zebrafish *angpt1* promotes cardiovascular activity and neurogenesis. (A) Overexpression of zebrafish *angpt1* by the ubiquitin promoter is revealed by WISH in 3-dpf larvae. Upregulation of *notch1a* is found in the proliferation zones of the forebrain ventricular zone and posterior midbrain in *angpt1* overexpressing zebrafish. (B) A higher heartbeat rate is found in the transgenic *angpt1* overexpressing larvae. (C) Microangiography of the vascular formation shows the disorganization of trunk blood vessels in the *angpt1* overexpressing larvae. (D) Quantification of the blood flow shows no significant differences. (E) Quantification of related mRNA expression levels in control and *angpt1* overexpressing groups. Data represent the mean ± SEM of three independent experiments (N=3 in each group and 10-pooled embryos in each group). *p<0.05 and ***p<0.001 by Student’s *t* test.

### Neural overexpression of zebrafish *angpt1* upregulates proliferation and HuC/D positive neuronal precursor cells in zebrafish larval brains

To observe whether ubiquitous, neural or non-neural overexpression of *angpt1* could affect embryonic neurogenesis, four transgenic constructs including zebrafish *angpt1* driven by the ubiquitin promoter, the *h2afx* promoter, the *elavl3* promoter, and the *gfap* promoter, were injected in Turku WT embryos. 7-dpf dissected brains with conditionally overexpressed *angpt1* were co-stained with antibodies recognizing GABA (a GABAergic marker), and HuC/D (a pan-neuronal marker) following the EdU staining (a proliferation marker). The positively labeled cells stained with these markers in the caudal hypothalamus (Hc) were quantified. We found a robust increase in proliferating cell numbers in the *elavl3*-driven *angpt1* group (Fig. 13G,13M) compared with those of the control-injected, the *h2afx*-driven, and the *gfap*-driven *angpt1* groups (Fig. 13A, 13D, and 13J). In contrast to a significant increase in proliferation, a substantial decrease in GABA-positive cell numbers was found in the *elavl3*-driven *angpt1* (Fig. 13H, 13N) and in the *gfap*-driven *angpt1* groups (Fig. 13K, 13N) compared with the control-injected and the *h2afx*-driven *angpt1* groups (Fig. 13B, 13E). A significant increase in HuC-positive cell numbers was found in the *h2afx* - driven (Fig. 13F) and the *elavl3*-driven *angpt1* groups (Fig. 13I, 13O) compared with the control-injected (Fig. 13C) and the *gfap*-driven *angpt1* groups (Fig. 13L). These data indicate that overexpression of *angpt1* in neuronal progenitor cells stimulates proliferation and differentiation but has a downregulating effect on GABAergic neurons in the caudal hypothalamus area.

**Fig. 13.**
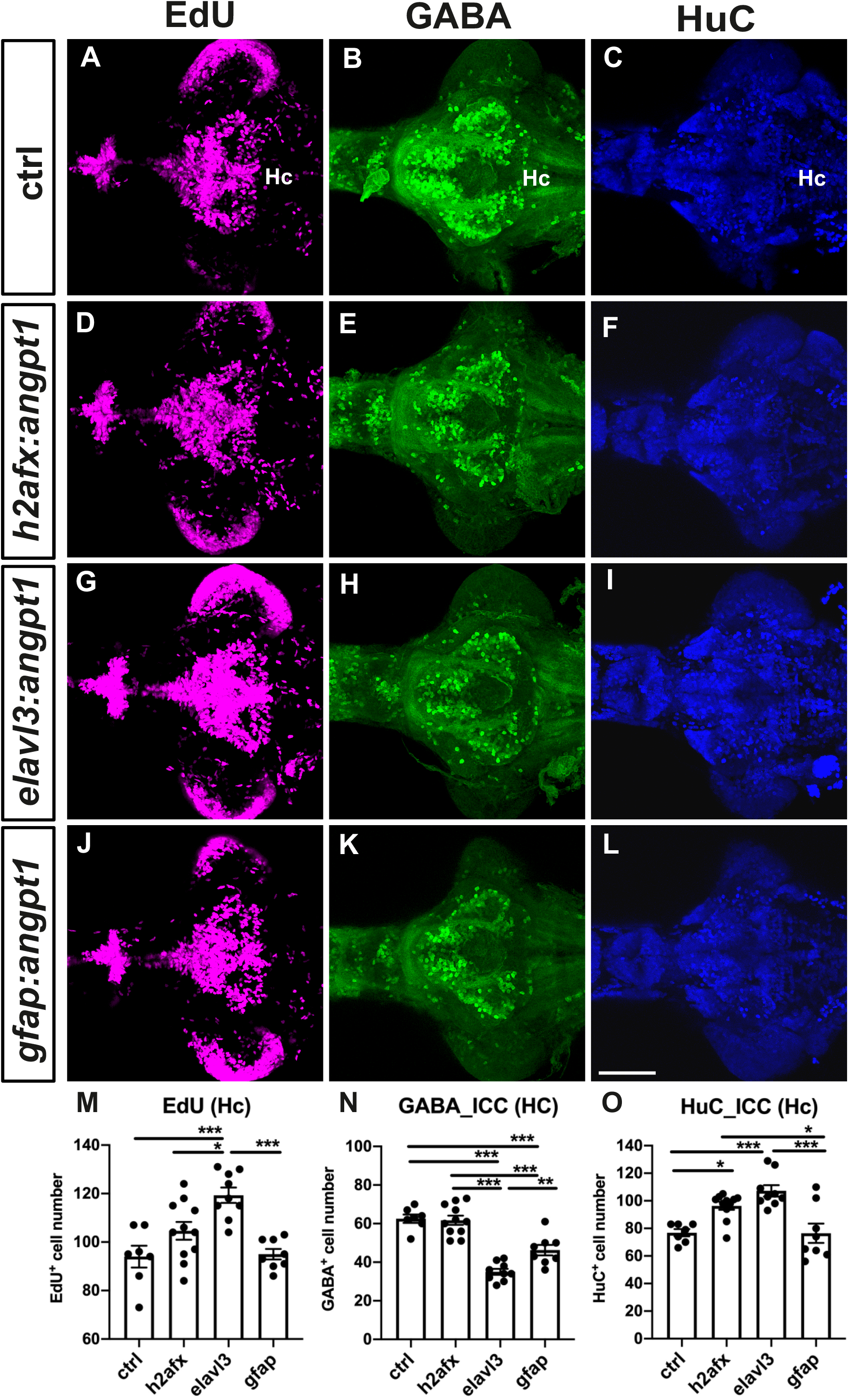
Ectopic expression of zebrafish *angpt1* upregulates proliferation and HuC/D positive neuronal precursor cells in 7-dpf zebrafish larval brains. Control-injected brains stained with (A) EdU, (B) GABA and (C) HuC. Larval brains with injected *h2afx* promoter driving *angpt1* stained with (D) EdU, (E) GABA and (F) HuC/D. Larval brains with injected *elavl3* promoter driving *angpt1* stained with (G) EdU, (H) GABA, and (I) HuC/D. Larval brains with injected *gfap* promoter driving angpt1 stained with (J) EdU, (K) GABA, and (L) HuC/D. (M) Quantification of EdU-labeled cell numbers in (the caudal hypothalamus (Hc). (N) Quantification of GABA immunoreactive cell numbers in the Hc area. (O) Quantification of HC/D-positive cells in the Hc area. Data are mean ± SEM. One-way ANOVA analysis with multiple comparisons was used for statistical analysis *p<0.05, **p<0.01, and ***p<0.001. Scale bar is 100μm.

### Neural overexpression of zebrafish *angpt1* promotes proliferation and neuronal precursors in *tek* mutant brains

Overexpression of *angpt1* stimulated the proliferation and differentiation in the Turku WT larval brains. To investigate whether *angpt1* and its primarily binding receptor *tek* effectively contributed to the regulation of the embryogenic neurogenesis, the *tek* mutant fish was subjected to overexpression of *angpt1* driven by the *elavl3* promoter to quantify the proliferating and neuronal cell numbers using immunostaining. A significant increase of the proliferating cells was found in the *elavl3*-driven *angpt1* injected *tek* ^+/+^ and *tek*^−/−^ groups (Fig. 14G, 14J and 14M) compared with the control-injected groups (Fig. 14A, 14D). The *elavl3*-driven *angpt1* injected *tek*^+/+^ group produced a more substantial effect on proliferation than in the *elavl3*-driven *angpt1 tek*^−/−^ group (Fig. 14M) whereas a significant reduction in GABAergic neurons was found in the *elavl3*-driven *angpt1 tek* ^+/+^ (Fig. 14H) and *tek*^−/−^ groups (Fig. 14K) compared with their control-injected groups (Fig. 14B, 14E and 14N). Moreover, in comparison with the control-injected groups (Fig. 14C, 14F), the number of HuC-positive cells was significantly increased (Fig. 14O) in the *elavl3*-driven *angpt1* injected *tek* ^+/+^ (Fig. 14I) and *tek*^−/−^ (Fig. 14L) groups. Our data indicate that *angpt1* stand-alone without the functional *tek* receptor still remains a significant impact on modulating embryonic neurogenesis in the zebrafish brain.

**Fig. 14.**
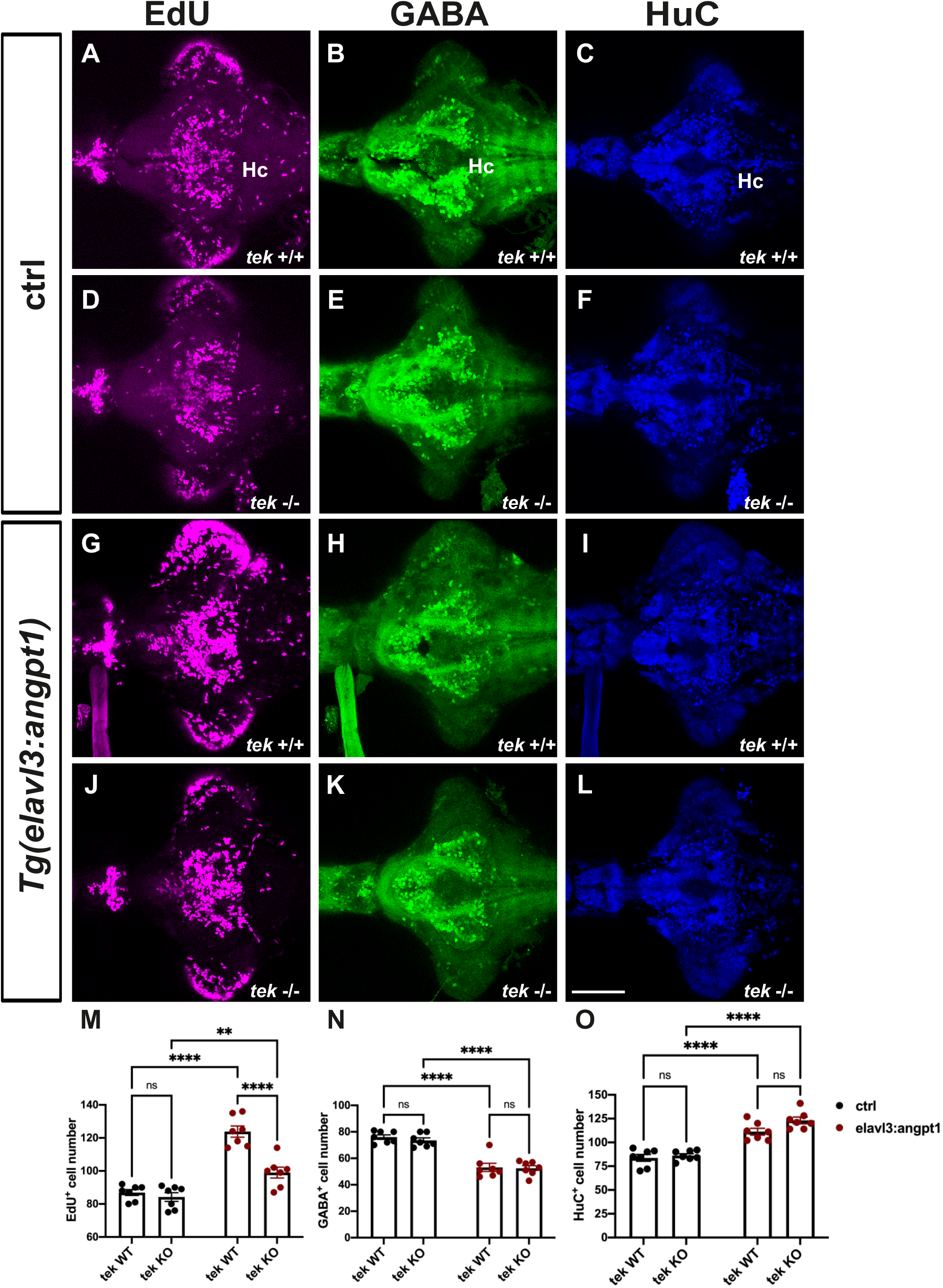
Neurally driven overexpression of zebrafish *angpt1* increases proliferation and neuronal precursors in *tek* mutant brains. 7-dpf *tek* WT and KO brains with *angpt1* overexpression driven by the *elavl3* promoter were triple labeled with EdU, anti-GABA, and anti-HuC/D antibodies. Control injected *tek* WT brains stained with (A) EdU, (B) GABA, and (C) HuC. Control injected *tek* KO brains stained with (D)EdU, (E) GABA, and (F) HuC. Neural overexpression of *tek* WT brains stained with (G) EdU, (H) GABA, and (I) HuC. Neural overexpression of tek KO brains stained with (J) EdU, (K) GABA, and (L) HuC. (M) Quantification of EdU-labled cell numbers in (the caudal hypothalamus (Hc). (N) Quantification of GABA immunoreactive cell numbers in the Hc area. (O) Quantification of HC/D-positive cells in the Hc area. Data are mean ± SEM. Two-way ANOVA analysis with multiple comparisons was used for statistical analysis **p<0.01, ***p<0.001 and ****p<0.0001. Scale bar is 100μm.

## Discussion

Our present data show for the first-time *in vivo* evidence that *angpt1* and *itgb1b* play not only essential roles in developing cerebrovascular formation but exert also own neurogenic effects on the development of different neural cell populations, including dopaminergic, histaminergic, and GABAergic systems, and the patterning of hindbrain reticulospinal neurons in the CNS. Our novel findings also provide evidence that the role of *tek* is dispensable in embryonic neurogenesis in zebrafish, similar to the published results in angiogenesis (Gjini et al., 2011; Jiang et al., 2020).

Besides indispensable roles in mammalian developmental cardiovascular formation and the maintenance of adult vascular stability, increasing evidence shows that after brain trauma, Angpt1 functions in a dual way in promoting angiogenesis and neurogenesis to enhance BBB integrity and neurological regeneration (Meng et al., 2014; Michinaga and Koyama, 2019; Wang et al., 2019). Moreover, in neural cell cultures, Angpt1 acts directly on neurite outgrowth via a beta1-integrin-dependent manner instead of binding to its typical receptor, Tek (Carlson et al., 2001; Chen et al., 2009a). However, the molecular mechanisms of neurogenic effects of *angpt1* and *itgb1b* in normal neurodevelopment remains mostly unrevealed. The Angpt1 and Tek knockout mice die between E9.5 and E12.5 because of cardiovascular failure (Sato et al., 1995; Suri et al., 1996). Therefore, embryonic lethality makes it impossible to observe the role of *Angpt1* and *Tek* in embryonic neurogenesis in mammals.

In this study, we take advantage of zebrafish fast *ex utero* development and individual neurotransmitter systems in the CNS already developed within 7dpf (Panula et al., 2010), which allows us to characterize the overt vascular formation and the evident neural phenotypes in *angpt1*, *itgb1b*, and *tek* mutant brains during embryonic development. However, the *angpt1* and *itgb1b* mutant fish exhibited lethality between 5 and 7 dpf. Our gain-of-function approach also demonstrated the neurogenic effect of zebrafish *angpt1* on promoting proliferation and differentiation in the developing brain. Moreover, in transgenic fish overexpressing angpt1, we observed disorganized vessel branches in the brain and the trunk, suggesting the importance of the precise spatiotemporal distribution of angpt1 in controlling vasculogenesis and angiogenesis. The angpt1 mutant larvae revealed severe deficiencies of angiogenesis in central arteries (CtAs) in the mid-and hindbrain; in contrast, the structure of the intersegmental vessels (Ses) in the trunk was not affected. The unaffected Se structure in the *angpt1* mutant larval trunk may be contributed by the upregulations of other angiogenic factors, including *itgb1b*, *angpt2*, and *vegfaa*, to compensate for the loss of *angpt1* (Felcht et al., 2012; Wild et al., 2017). Additionally, upregulated *angpt2a* could play a dual role in compensation of neuronal (Liu et al., 2009) and vascular defects. Although Angpt2 mainly recognizes as an antagonistic ligand of Tek, it has been reported to function as an agonist in the absence of Angpt1 (Daly et al., 2013).

In embryonic neurogenesis, the Notch1 signaling pathway plays a crucial role in maintaining NSC proliferation and promoting radial precursor cell fates whereas restricting NSC differentiation (Koch et al., 2013). In periphery, NOTCH receptors and their ligands take a major part in regulation of cardiomyocyte proliferation and arterial-venous determination (Lin et al., 2019). In our previous study, an upregulated *notch1* pattern was observed in the 2- and 6-dpf brains of zebrafish *angpt1* knockdown morphants (Chen et al., 2015). As opposed to this morphant result, downregulation of *notch1* transcripts was observed in the *angpt1*^*−/−*^ embryos. The discrepancy in the *notch1* expression in the brain between morphants and null mutant embryos may be primarily due to the activation of genetic compensation in the *angpt1*^*−/−*^ embryos, a compensatory effect that unlikely occurs in the transient morpholino knockdown approach (El-Brolosy et al., 2019; Rossi et al., 2015), although an unexpected effect of morpholinos cannot be entirely ruled out either. Thus, both approaches showed reproducible phenotypes: a smaller brain and embryonic lethality when angpt1 is deficient.

### Angpt1 and itgb1b regulate neural proliferation during neurogenesis

During embryonic neurogenesis, to drive neuroepithelial cells from the proliferating state to mature neurons, neuronal stem cells (NSCs) vitally require the vasculature to deliver oxygen, nutrients, and specific signaling cues to regulate the fate of NSCs in a timely and accurate manner (Tata et al., 2015; Walchli et al., 2015). Moreover, NSCs need adequate oxygenation from the nearby microvasculature to restrain hypoxia stress to switch from the self-renewal radial glial stem cells to neuronal progenitors (Lange et al., 2016). When low oxygen tension occurs in neurogenesis, the expansion of radial glial cells pauses at the symmetrical self-renewal state and fails to commit the subsequent steps (Schmidt et al., 2013). This disturbance will restrict the number of neuron progenitors promoting to mature neurons, ultimately resulting in a small-sized brain. For instance, G protein-coupled receptor 124 (GPR124; tumor endothelial marker 5, TEM5) is required for the CNS vascularization and BBB formation. GPR124-deficient mice appear to have a prominently disorganized vasculature, decreased cortical thickness, and reduced brain volume (Lange et al., 2016). Similarly, in the angpt1 (this study) and itgb1a-deficient zebrafish (Iida et al., 2018), the cerebrovasculature was severely impaired during development. Parallelly, the deficiency of proliferation but a robust increase of the gfap-positive radial glia cells in the hindbrain was found in these two mutant brains, which were accompanied by a significant down-regulation of neurogenic genes including *pcna, notch1a, nestin, wnt1*, and *sox2*. Subsequently, the deficient neurogenesis resulted in a considerable decrease of dopaminergic and histaminergic neurons in the *angpt1* and *itgb1b* mutant brain. Therefore, insufficient oxygen supply causes hypoxia-induced radial glia expansion. As a result, it produces a neurogenic disruption from the proliferation to neuronal differentiation, consequently reducing the brain size that may explain the cause of brain defects in these two mutants. However, in zebrafish, embryonic neurogenesis starts at 10 hpf (early somitogenesis) (Schmidt et al., 2013), earlier than cerebral vessel sprouting at around 24 hpf (late somitogenesis)(Ulrich et al., 2011). Additionally, zebrafish angpt1 was detectable starting from the maternal stage, indicating that angpt1 may act as a potent pro-neurogenic factor in regulating embryonic neurogenesis prior to its role in angiogenesis. Consistent with this current finding, in the guppy, the larger brain population has the higher expression level of *angpt1* compared with the smaller brain population (Chen et al., 2015).

### Angpt1 affects the maturation of brain dopaminergic, histaminergic and GABAergic neurons

Lack of *angpt1* caused the dopaminergic and histaminergic neuron deficiency mainly due to the impaired neurogenesis. Notably, ectopically overexpressed zebrafish angpt1 in pan-neuronal cells promoted a dramatic increase of proliferation and neuronal progenitors; in contrast, a decreased number of GABAergic neurons was found in these transgenic larval brains. During neurogenesis, the nascent vasculature offers not only oxygen and certain signaling factors to drive neural proliferation and differentiation (Li et al., 2018; Tan et al., 2016), but it also serves as a guiding framework for newly born neurons migrating to the right locations (Won et al., 2013). Our gain-of-function models caused a deficiency of GABAergic neurons may be due to two aspects: one possibility is that disorganized angiogenesis misleads GABAergic progenitors to the wrong targets and trigger neural apoptosis resulting in a reduction of GABAergic neurons (Licht and Keshet, 2015); on the other hand, ectopic expression of angpt1 associated with the extensive proliferation may negatively control the maturation of GABAergic progenitor neurons resulting in a decrease of the GABAergic neurons although we have no evidence to prove that *angpt1* is directly involved in regulation of GABAergic development.

### Angpt1 and itgb1b are required for hindbrain neurovascular development

The hindbrain organization requires interdependence between the genetic, morphological, and neuroanatomical segmentation because a disordered morphological segmentation will disrupt the hindbrain neuron organizations and vice versa (Moens and Prince, 2002). In zebrafish brain vascularization, the primordial hindbrain channels (PHBCs) arise at around 20 hpf and extend laterally along the anterior-posterior axis and later contribute to hindbrain vascularization. The first cerebral angiogenic sprouts developing from the inner lateral side of both PHBCs integrate correspondingly into neuroepithelium alongside the rhombomere boundaries after 28 hpf (Fujita et al., 2011; Ulrich et al., 2011). In zebrafish hindbrain organization, Mauthner cells are the largest reticulospinal neurons, reside bilaterally in rhombomere 4 (r4), and are the first identified hindbrain neurons beginning at 8-10 hpf (during gastrulation). No later than 18 hpf, axonal projections penetrate the segments in a stereotyped manner before the neuromeres are formed at 18-20 hpf (Moens and Prince, 2002). Accordingly, the branchiomotor (BM) neurons of the cranial nerves and the reticulospinal (RS) interneurons develop in a rhombomere-specific disposition and provide markers of segmental identity (Moens and Prince, 2002). The specificity of each segment and the identity of rhombomere boundaries requires transcription factors, angioneurins, and Wnt-delta-Notch signaling pathways to maintain rhombomeres’ specification (Riley et al., 2004). For instance, Krox-20, a zinc finger transcription factor is required to develop rhombomeres 3 and 5 and maintain regional specification. Loss of functional Krox-20 causes disorganized odd- and even-numbered territories alongside the hindbrain and neuronal organization corresponding to the rhombomeres (Schneider-Maunoury et al., 1993; Voiculescu et al., 2001). Angioneurins, secreted and transmembrane proteins such as VEGF and Eph-ephrin, empower both vasculotrophic and neurotrophic activities (Schwarz et al., 2004; Xu et al., 2000). Impairment of these factors disrupts the regular segment-restricted expression of krox-20 in the zebrafish hindbrain (Giudicelli et al., 2001; Laussu et al., 2017). Wnt1 regulates neurogenesis and mediates lateral inhibition of boundary cell specification in the zebrafish hindbrain via Notch signaling pathways (Amoyel et al., 2005; Cooke et al., 2001; Xu et al., 1995). Since the neural organization occurs earlier than the vessel invasions in the hindbrain formation, in the *angpt1* and *itgb1b* mutant hindbrain, the deficiency of reticulospinal neurons accompanied with downregulation of *notch1a, wnt1* and *krox20* renders it possible that angpt1 and itgb1b may function in angioneurin-like manners having neurotrophic effects regulating segment-specific fates and specifying rhombomeric territories even before angiogenesis proceeds.

### In summary

In this study, we used both loss-of-function and gain-of-function approaches to provide novel evidence that angpt1 and itgb1b perform dual functions in zebrafish angiogenesis as well as in embryonic neurogenesis for developing normal hindbrain morphogenesis and particularly, promoting neural proliferation in a *tek*-independent manner. Although this study did not demonstrate the interaction between angpt1 and itgb1b as the previous study shown overexpression of α5β1 integrins and angpt1 stimulating angiogenesis following ischemic stroke (Wang et al., 2019). Our findings highlight the important-neurogenic effects of angpt1 and its putative receptor itgb1b that may support the concept of angiopoietin-based treatments for the clinical therapeutics in neurological disorders such as Alzheimer’s disease, stroke and traumatic brain injuries (Venkat et al., 2021; Zlokovic, 2011).

## Acknowledgements

This study was supported by the Jane and Aatos Erkko Foundation, Finska Läkaresällskapet, and Magnus Ehrnrooth’s Foundation. We thank Dr. Stefan Schulte-Merker for the *tek*^*hu1667*^ mutant line and Dr. Koichi Kawakami for the Tol2 plasmid. Henri Koivula, BSc, Niina Siiskonen, BSc, and Sanni Perttunen, BSc, are thanked for expert technical help. Zebrafish used in this study were produced and maintained at the Zebrafish Unit of the HiLife infrastructure of the University of Helsinki, supported by Biocenter Finland.

## Author contributions

Y.-C. C. and P.P. designed the study and interpreted results, Y.-C.C., T.A.M. and V.M. carried out experiments, Y.C. and P.P. wrote the paper, all authors commented and revised the paper.

